# Discontinuities in quinoa biodiversity in the dry Andes: an 18-century perspective based on allelic genotyping

**DOI:** 10.1101/366286

**Authors:** Thierry Winkel, María Gabriela Aguirre, Carla Marcela Arizio, Carlos Alberto Aschero, María del Pilar Babot, Laure Benoit, Concetta Burgarella, Sabrina Costa-Tártara, Marie-Pierre Dubois, Laurène Gay, Salomón Hocsman, Margaux Jullien, Sara María Luisa López-Campeny, María Marcela Manifesto, Miguel Navascués, Nurit Oliszewski, Elizabeth Pintar, Saliha Zenboudji, Héctor Daniel Bertero, Richard Joffre

## Abstract

History and environment shape crop biodiversity, particularly in areas with vulnerable human communities and ecosystems. Tracing crop biodiversity over time helps understand how rural societies cope with anthropogenic or climatic changes. Exceptionally well preserved ancient DNA of quinoa (*Chenopodium quinoa* Willd.) from the cold and arid Andes of Argentina has allowed us to track changes and continuities in quinoa diversity over 18 centuries, by coupling genotyping of 157 ancient and modern seeds by 24 SSR markers with cluster and coalescence analyses. Cluster analyses revealed clear population patterns separating modern and ancient quinoas. Coalescence-based analyses revealed that genetic drift within a single population cannot explain genetic differentiation among ancient and modern quinoas. The hypothesis of a genetic bottleneck related to the Spanish Conquest also does not seem to apply at a local scale. Instead, the most likely scenario is the replacement of preexisting quinoa gene pools with new ones of lower genetic diversity. This process occurred at least twice in the last 18 centuries: first, between the 6th and 12th centuries—a time of agricultural intensification well before the Inka and Spanish conquests—and then between the 13th century and today—a period marked by farming marginalization in the late 19th century likely due to a severe multidecadal drought. While these processes of local gene pool replacement do not imply losses of genetic diversity at the metapopulation scale, they support the view that gene pool replacement linked to social and environmental changes can result from opposite agricultural trajectories.

## Introduction

The Andes, a global hotspot of past and present crop biodiversity, has witnessed huge environmental and socio-cultural changes, including the climatic fluctuations of the late Holocene and the disruption of native societies following the Spanish Conquest (1-3). Less dramatically, progressive changes in agricultural knowledge and practices have ensured the resilience of Andean societies to date (4-6). Amid these historical changes, several Andean-origin crops have diversified and were successfully disseminated throughout the world, such as tomato (*Solanum lycopersicum*), potato (S. *tuberosum*), beans (*Phaseolus* spp.), chiles (*Capsicum* spp.) and, more recently, quinoa (*Chenopodium quinoa*) (7).

In the Central Andes of Peru and northern Bolivia, the rise and fall of past agrarian societies due to political and environmental changes seems a most likely scenario (8-10). But in the dry Andes of Northwest Argentina, southern Bolivia and northern Chile (**Fig 1A**), a different historical trajectory took place due to the relative importance of pastoralism versus agriculture (11) (**SI-1 Table**). Around 5000 BP (years before present) pastoralism arose among local hunter-gatherers who had been established in the region since 12000 BP (12, 13). These hunter-gatherers in transition to food production also developed crop planting early in the dry Andes, as evidenced by remains of plant domesticates dating back *ca* 5000 BP (14, 15). Farming was a productive practice in the region at that time and until the Inka period and the Spanish conquest, though without reaching a comparable level of significance to that observed in the Central Andes (16, 17). Then, at a still uncertain time during the Colonia and early Republic periods (*viz*. 16th to 19th centuries), agrarian systems in the most arid highlands reverted to a primarily pastoralist economy, wherein small-scale crop farming assumed a limited role, a situation that persists today (18). Palaeoecological studies revealed that substantial Holocene fluctuations in the regional climate likely coincided with these socio-historical changes (19-22). Two relatively humid periods (12000-8000 BP and 5000-1500 BP) alternated with drier periods (8000-5000 BP and 1500 BP to the Present); during the dry phases water resources concentrated in some valleys and basins (23).

**Fig 1.**
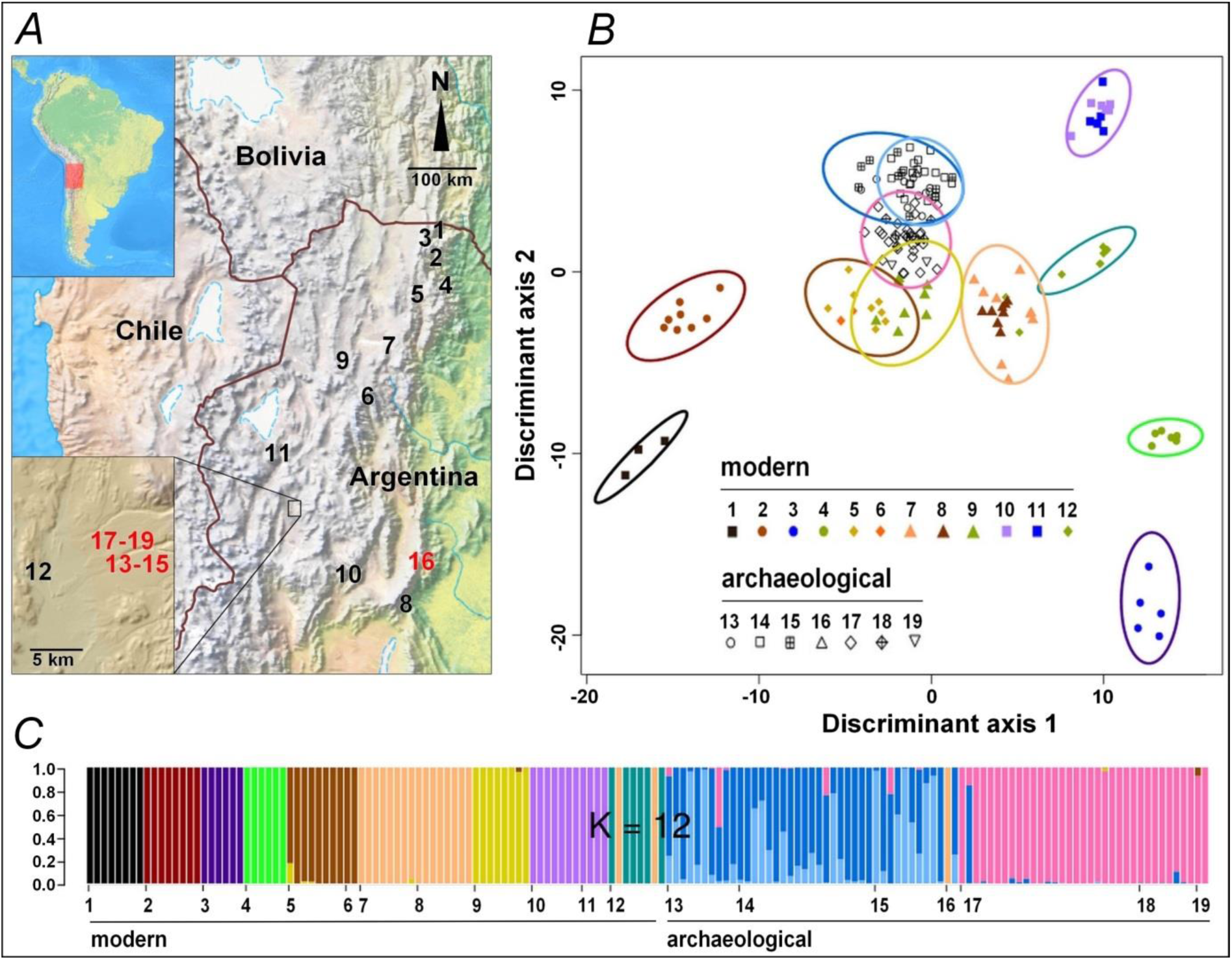
Geographic localization and genetic classification of ancient and modern quinoas collected in Northwest Argentina. (*A*): Map of the dry Andes localizing ancient and modern quinoa samples (red and black numbers, respectively; detailed sample description in **SI-2 Table**). (*B*): Scatterplot of the Discriminant Analysis of Principal Components (DAPC). Individuals are represented by symbols according to their sample of origin; colored inertia ellipses define the clusters identified with a k-means algorithm for k=12. (*C*): Individual assignment probability to each cluster from DAPC. Horizontal axis shows the sample codes as in *A* and *B*; vertical axis shows the assignment probabilities for k=12.

The question then arises as to how these social and environmental changes affected local crop biodiversity throughout this period. Specifically, have climatic and agrarian changes—and their related transformations in social structures and local economy—led to genetic changes in native crop species? The quinoa crop in the dry Andes of Argentina provides a case study for investigating these issues since local conditions of low temperature and air dryness allowed for the conservation of abundant biological material in what once were residential places, granaries or tombs (24, 25). In modern quinoa samples, molecular markers reveal a diversity essentially shaped by broad biogeographic features separating—among others—quinoas from temperate highlands, arid highlands, mid- and high-altitude valleys, and western versus eastern lowlands (26). Molecular genotyping, applied to ancient samples, should thus allow for tracking quinoa biodiversity in space and time, providing a new tool to investigate the agrarian economy of past societies and go further in-depth into the history of human-plant relationships (27).

Analyzing genetic markers within a coalescence framework, we track changes and continuities in quinoa diversity in the dry Andes over the last two millennia. Coalescence theory allows to identify the most probable trajectory among the many possible genealogies in a regional gene pool (28). Then we discuss how natural and human circumstances paralleling these temporal patterns in genetic diversity could explain them. Our archaeological study sites are located in cold and arid highlands, with one site in a mesothermic Andean valley located at the same latitude (**Fig 1A, SI-2 Table**) (12, 14, 17, 29). They provided well-preserved quinoa seeds, with a broad chronological range spanning the time of early husbandry (*ca* 1800 BP), to periods of stable agro-pastoralist societies (*ca* 1400 BP) and complex corporative societies (*ca* 800-700 BP). To evaluate the relationship of these ancient quinoas with the present-day germplasm, we studied a reference panel of quinoas collected in 2006-2007 from different environments in the Andean highlands of Argentina (26, 30) (**Fig 1A, SI-2 Table**). Some archaeological sites supplied both dark and white seeds, which allowed us to explore the diversity of cultivated quinoa (generally white-seeded) and their weedy relatives (all dark-seeded).

## Results

Genetic diversity, selfing rates and genetic structure were studied from 157 quinoa seeds (76 ancient, 81 modern) genotyped at 24 microsatellite loci (see **SI-3 Text** for detailed results).

### Genetic diversity and selfing rate

Estimates of the diversity indices and the selfing rates of the 19 populations sampled are summarized in table SI-3.3. Expected heterozygosity (He) in the subset of modern quinoas showed highly variable values (range 0.02-0.70), consistent with those found in an independent study on the same samples (26). Similarly, selfing rates (*s*(*F_is_*) and *s*(*LnL*)) appear highly variable without any clear geographical pattern. Comparing quinoa samples through time at Antofagasta de la Sierra (hereafter: Antofagasta) shows a trend towards lower allelic diversity (*N_all-rar_*) and expected heterozygosity (*H_e_*) in the modern sample (#12) compared to the ancient ones (#13-15, 17-19). Selfing rate (*s(LnL)*) increased significantly (P<0.05) from the most ancient samples (#17-18) to the intermediate (#13-14) and modern ones (#12).

### Genetic structure in time and space

The discriminant analysis reveals a neat distinction between ancient and modern samples, ancient samples showing little affinity to modern samples, particularly for the geographically closer sample #12 (**Fig 1B, SI-3.4 Table**). Among modern samples, genetic structure reflected the geographical sampling, with most samples showing a marked identity (**Figs 1B-C, Figs SI-3.4, SI-3.5**). Some sites showed strong affinities between them with a clear ecogeographical link (samples #1,2 are from NE humid valleys, and samples #10,11 from arid highlands) while others are more difficult to interpret in simple ecogeographical terms (sample #8, from a mid-altitude mesothermic valley, shows strong affinity to sample #7, from cold and arid highlands).

Among ancient samples, genetic structure is associated to the age of the seeds. The two main clusters identified separate the more recent samples (#13–15) from the older samples (#17–19) (**Fig 1B, Figs SI-3.4, SI-3.5**). Affinities of sample #16 are unclear as it is composed of only two seeds with very different genotypes and from a geographically distant location. Although collected from residential or storage places, or from ritual deposits, all likely to have received seeds from various fields or sites, the ancient samples (#13-19) each showed a fairly high homogeneity, similar to that of modern samples collected in separate fields. Samples #13-15 (*ca* 690-796 cal BP) grouped together in spite of differences in seed color (#13,14 are white, #15 is dark). The two seeds from sample #16 (*ca* 1270 cal BP) were assigned to this same group. Samples #17 and 18 (*ca* 1364 cal BP) were assigned to a distinct group, which also included the oldest sample #19 (*ca* 1796 cal BP), being differently colored (#17,19 are white, #18 is dark). Interestingly, additional structure analysis separates dark-seeded quinoa samples (#15,18) and white-seeded samples from the same age and locality (#14,17 respectively) (**Fig SI-3.5**, k=12). Yet the wild-form samples remain genetically close to the white-seeded samples of their respective time x location set.

### Inference of local demographic history

At Antofagasta, one modern sample and six ancient samples offer a temporal series covering almost two millennia. Analyses of genetic structure of these samples reveal three main genetic groups: modern (#12), intermediate (#13–15), and ancient (#17–19) (**Fig 1C**). The differentiation among them could be due to genetic drift between sampling times within a single population or to divergence among distinct populations that locally arrive at different times, replacing the preexisting genetic pool. In order to distinguish between these two alternative processes we built demographic scenarios in which the two temporal and genetic discontinuities are either simulated as genetic drift within a single population (**Fig 2A**), or mixtures of drift and replacement (**Fig 2B-D**). As dark seed quinoa samples (#15,18) show some differentiation from white seeds (see **Fig SI-3.4** for k=14 and **Fig SI-3.5** for k=9 or higher), two additional scenarios were tested: one differentiating between cultivated and wild compartments (**Fig 2E**), the other considering an admixture event between both compartments (**Fig 2F**).

**Fig 2.**
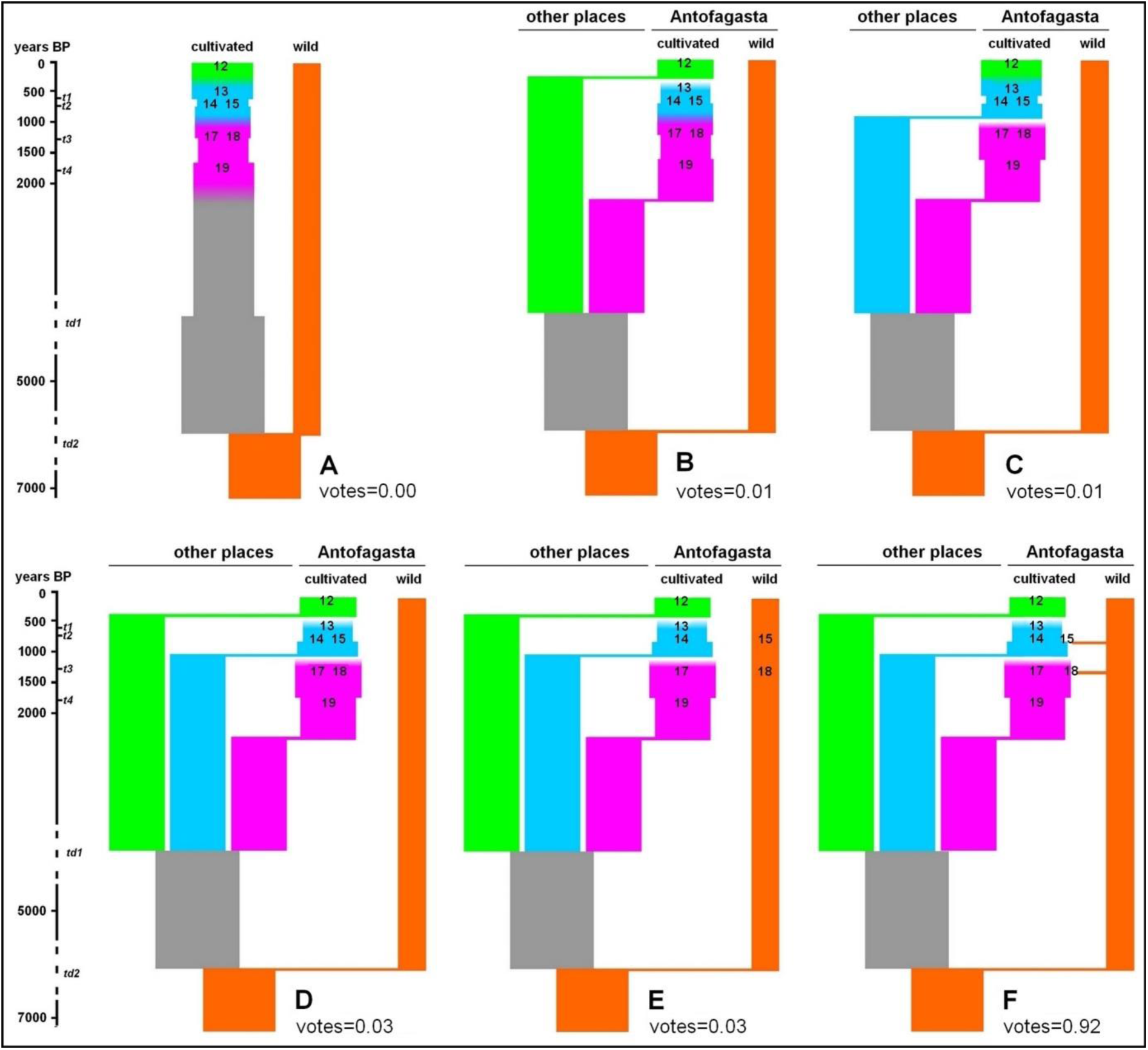
Graphical representation of alternative genetic relationships between modern and ancient quinoas from Antofagasta. The coalescence-based modeling examined six scenarios: (*A*) direct chronological filiation between all samples, mixing cultivated and wild forms, (*B*) replacement of all ancient quinoas by modern quinoas, (*C*) filiation between modern quinoas and intermediate ancient quinoas, both replacing the oldest quinoas, (*D*) successive replacement of the three groups of quinoas: modern, intermediate, ancient, (*E*) same scenario as previously but differentiating between cultivated and wild forms, (*F*) same scenario as previously but with admixture between cultivated and wild forms. Numbers in bold refer to modern (#12) and ancient quinoa samples (#13-15,17-19) described in **SI-2 Table**. Vertical axis shows years BP with present at the top. Proportion of votes received from the random forest classification of the observed data is shown for each model. *t_n_* and *td_n_* are time parameters used in the approximate Bayesian computation analysis (see **SI-3.5 Table**).

The coalescence analysis clearly identifies as the model with the best fit (votes=0.92, posterior probability=0.99,) the demographic scenario in which three genetic clusters belong to three separate cultivated gene pools, with gene flow from a wild compartment producing dark seeds (**Fig 2F**). We estimated effective population sizes for the 3 quinoa samples around few 100 individuals and around 30 individuals for the wild gene pool (**SI-3.5 Table)**. Admixture proportions from the wild pool of dark seeds was high but lower than 0.5 (point estimates and 95% highest posterior density intervals are reported in **SI-3.5 Table**). Posterior probability distributions for other parameters of the model (e.g. ancestral population effective size, time of divergence) were indistinguishable from priors (**SI-3.5 Table**).

## Discussion

Using a coalescent-based approach, this study provides the first evidence of a significant change in the demographic history of quinoa in the Andes over 18 centuries. The analysis of modern and ancient samples from Antofagasta reveals that genetic differentiation among samples from different times cannot be explained by genetic drift. Instead, the most likely scenario in this locality is the replacement of preexisting quinoa gene pools with new exogenous gene pool. This process occurred at least twice in the last 18 centuries: first, between 1364 and 796 cal BP, well before the Inka and Spanish conquests—respectively initiated 568 and 483 years ago in Northwest Argentina (31)—, and then between 690 cal BP and today, an interval of time covering the Inka, Colonial and Republican periods. The general assumption of successive genetic bottlenecks for the Andean quinoa—related to the initial events of hybridization and domestication, and then to the Spanish Conquest (32)—thus does not seem to apply at a local scale. This should be tracked back now over a larger geographic area, particularly the Central Andes where less extreme climatic conditions and a distinct socio-historical context might have led to other patterns of genetic change in quinoa. In the dry Andes, these two events of gene pool replacement appeared associated with quite different socio-environmental dynamics, namely: a phase of *agricultural intensification* initiated 1100 years ago followed by an opposite phase of *farming marginalization* in the Colonial and Republican periods.

### Intensification of agriculture

Intensification of agriculture by local societies starting 1100 BP was contemporary to the increasing aridity, which reduced water availability for crops and pastures in the region (13, 17). Large irrigation infrastructures were then established, probably associated with an increase in population density (29). Although rare in the region, these intensified crop-pasture systems allowed to expand the productive land area from small humid river banks to broader alluvial terraces (33). Adaptation of quinoa to these new climate and farming conditions could have occurred in two ways: either by local selection for ever more drought-tolerant variants or variants apt for new intensified fields, or directly by replacing local varieties with new ones from other regions. Our model of quinoa demographic history in Antofagasta showing the introduction of a new gene pool in the 1364-796 BP period (**Fig 2F**) is congruent with the second hypothesis, and concurs with the intense interregional connections at that time (29). The grouping of samples #13-15 from Antofagasta with sample #16 from mesothermal valleys (**Figs 1B-C**) suggests that the same gene pool might have circulated between dry highlands and valleys in the 1270-690 BP period.

As reported in other Andean regions (34), modern weedy (dark-seeded) quinoa appeared genetically related to sympatric cultivated (white-seeded) quinoa populations. Black chenopod seeds are generally assigned to the weed sub-species *Ch. quinoa* ssp. *melanospermum* and their relative frequency in archaeological remains is indicative of the degree of seed selection by past cultivators (35, 36). The presence of dark seeds (#15,18) in archaeological food processing places suggests the prolonged use of a combination of domesticated and weed chenopod grains by past populations in Northwest Argentina (37), a feature also observed elsewhere in the Americas (36, 38).

Another likely cause of the changes in quinoa demographic history relates to evidence of a generalized warfare in the dry Andes in the 750-600 BP period (11, 39, 40), a situation exacerbated by the competition for scarce water resources (11, 40), likely disturbing local seed-supply networks. Compared to the previous social system based on small villages, more complex and authority-centered societies at that time (41, 42) could also have impacted on seed availability and circulation in a trend towards less diverse crop practices and genetic resources.

In this context of coincident changes in climate, crop technology and society, our estimates of effective population size (**SI-3.5 Table**) suggest that the quinoa gene pool cultivated at Antofagasta *ca* 796-690 BP (samples #13,14) had a narrower genetic base and higher selfing rates than in the previous periods (samples #17,19). As the scenario of genetic drift within a single population is rejected by the coalescent-based analyses, the lower diversity of the cultivated quinoa at Antofagasta *ca* 796-690 BP is explained more by the displacement of local varieties by introduced ones with a narrower genetic base than by alternative hypotheses of enhanced selection for new cropping systems or loss of genetic resources due to endemic political unrest. In this perspective, agricultural intensification with newly introduced varieties can be considered as a risk-buffering strategy developed by ancient Andean peoples who, like other societies in the world, sought to ensure food security in a context of rising population, political conflicts and deteriorating climate (43, 44). This observation supports the idea that social and environmental stress can stimulate cultural innovation (4, 16). The brief Inka rule at this extreme end of the Andes continued this process of agricultural intensification as suggested by the appearance of large, albeit scattered, terrace and irrigation systems in the region (17, 45).

### Crop farming marginalization

Crop farming marginalization in the Andean highlands has been frequently attributed to the Spanish conquest (46-48). Undeniably, the European intrusion affected the structure of native societies and their subsistence systems across the Andes, including local farming activities (2, 49-51). Still, in the dry Andes the new mercantilist order prioritizing mining and caravan trading remained dependent on local crop-pasture systems for its food and forage supply (52, 53). Recent studies report the continuation, after the Spanish conquest, of local crop-pasture systems and food-storage facilities which allowed native populations in the remote highlands of Norwest Argentina to preserve relative autonomy and control over natural resources during the Colonial and Republican periods (33). Yet, these persisting crop-pasture systems stood vulnerable to climatic variations. A multidecadal drought in the 1860-1890s caused a severe mortality in the region (54), likely affecting local agriculture. Immediately after that time, socioecological changes related to emergent industrialization, urbanization, and globalization led to further rural depopulation and cropland abandonment throughout the 20th century (18). Under these cumulative factors, the relatively intensified crop-pasture systems built up during the pre-Hispanic period—and partly maintained until the 19th century—were dismantled in the study area and local agriculture returned to small-scale cropping and extensive pastoralism. In some southern highlands, intensified agricultural fields could have fallen into disuse much earlier—late 18th century or earlier—due to an emphasis on animal husbandry, whereas crop production continued as an important activity in the neighboring mesothermal valleys (50). We found that in the Colonial and Republican periods, a second event of gene pool replacement occurred in the quinoa cultivated at Antofagasta, which resulted in local quinoa gene pools of lower allelic diversity (**SI-3.3 Table**). As shown by population genetics theory (55), such a process of local gene pool replacement does not necessarily imply a loss of genetic diversity through time at the metapopulation scale. It does, however, support the view that gene pool replacement linked to social and environmental changes can result from opposite trajectories of agricultural intensification or marginalization. Such historical shifts in farming activities are characteristic of agriculture in extreme environments (37, 56, 57) and not only in remote times (58, 59).

## Material and methods

See the Supporting Information Appendix for an extended version of the methods.

### Seed sample collection, archaeological material and datings

Ancient and modern quinoa seed samples were collected from the sites described in **Fig 1A** and **SI-2 Table**. Archaeologists collected intact, non-charred samples of ancient quinoa in five sites related to agro-pastoralist societies from Northwest Argentina, covering the time span 1796-690 BP (detailed description in **SI-1 Text**). Four of the archaeological sites are located in arid highlands near the town of Antofagasta de la Sierra (Catamarca province): Cueva Salamanca 1 (sample #19; (60)), Punta de la Peña 9 (samples #17,18), Punta de la Peña E (samples #14,15), and Punta de la Peña 4 (sample #13; (61)). The fifth site, Cueva de los Corrales 1 (sample #16; (62)) corresponds to an area of mesothermal valleys in the Tucuman province. Sedimentary samples containing exceptionally preserved ancient quinoa remains from these sites were submitted to laboratory separation and concentration techniques (dry sieving and picking under magnifying glass) shortly before AMS dating and molecular analysis of seeds. In 2006-2007, an independent research team collected modern quinoa seed samples from 12 sites representative of different environments in the Andean highlands and valleys of Argentina (26, 30). Ancient and modern quinoa seeds were not in contact during their sampling, storage and manipulations.

### DNA extraction, microsatellite genotyping, and genetic data analysis

We extracted DNA from 81 modern and 144 ancient quinoa seeds according to the procedures described in **SI-3 Text**. DNA extraction was successful for all the modern seeds, while we recovered well-preserved DNA from only 76 ancient seeds (53%). To avoid contamination between ancient and modern DNA, we rigorously separated in time and space all the DNA extraction, DNA quality control and microsatellite amplification procedures detailed in **SI-3 Text**. We started by working on the ancient archaeological quinoas in a specific laboratory dedicated to ancient DNA. Once all the extractions and amplifications of ancient quinoa seeds were completed, we then proceeded to the extraction and amplification of modern quinoas in a distant laboratory, without any spatial connection with the previous one.

Ancient and modern quinoas were genotyped using 24 microsatellite loci (**SI-3.1 Table**). We did not find an ancient genotype identical to any other ancient or modern genotype, which proves the absence of contamination (see **SI-3 Text**). Descriptive genetic diversity (allelic richness, heterozygosity), inbreeding fixation coefficient (*F*_IS_) and genetic differentiation (*F*_ST_) were calculated in R using the packages *adegenet* and *hierfstat* (63, 64). The number of multilocus genotypes (*MLGs*) was computed in R using the package *poppr* (65). Diversity indexes were standardized using a rarefaction approach in ADZE (66). We estimated selfing rates for each sample in two independent ways, either from *F*_IS_, or using the maximum likelihood approach implemented in RMES (see **SI-3 Text**). The genetic structure of the samples was examined using the program STRUCTURE (67), principal component analysis (PCA), and discriminant analysis of principal components (DAPC) using *adegenet*. An approximate Bayesian computation approach using random forests (ABC-RF) (68, 69) was used to evaluate alternative models of demographic history of quinoa found around Antofagasta where one modern (#12) and six ancient seed samples (#13–15, #17–19) offer a temporal series covering 18 centuries. A classification vote system, which represents the frequency of each alternative model in the collection of classification trees, identified the model best suited to the observed dataset (68).

## Acknowledgements

All necessary permits were obtained for the described study, which complied with all relevant regulations. Archaeological field surveys received permits from the “Dirección de Patrimonio Cultural, Ente Cultural de Tucumán, Gobierno de la provincia de Tucumán, Argentina” and the “Dirección Provincial de Antropología, Gobierno de la Provincia de Catamarca, Argentina”. The export of ancient quinoa seeds for their analysis at CEFE received permit from the “Registro Nacional de Yacimientos, Colecciones y Objetos Arqueológicos, Instituto Nacional de Antropología y Pensamiento Latinoamericano, Argentina” (RENYCOA-INAPL). We thank Christelle Tougard and Claudine Montgelard for helpful advices in analyzing ancient DNA. We further thank Philippe Jarne and Eric Jellen for constructive comments on the manuscript.

## Supporting Information

### SI-1. ARCHAEOLOGICAL SITE DESCRIPTION

Four of the sites analyzed in this study are located in a dry and cold highland environment (*puna*) between 3600 and 3700 masl. They belong to the Punilla River basin in the Catamarca province (Argentina). The fifth site, Cueva de los Corrales 1, corresponds to an area of mesothermal valleys at 3000 masl in the Tucuman province. We present here the sites in the chronological order, starting from the most recent one.

The site **Punta de la Peña 4 (sample #13: AP4)** is a rock-shelter with a large protected area that occurs in the upper portion of the ignimbrite cliff of Punta de la Peña. Its archaeological occupation dates back to *ca* 9000 Before Present (BP). The layers 1 to 3 dated between *ca* 760 and 460 cal BP, with a sandy-silty sedimentary matrix. A high density and diversity of archaeological remains in the form of artifacts, ecofacts and structures with well-preserved plant and faunal remains characterized these layers. Within layer 3, the 3x lens was composed of a pit fire associated to numerous intact (dried) or charred quinoa seeds dated *ca* 690 ± 50 cal BP (UGA-15090, cal. 2σ, 95.4%: 1228-1398 Common Era (CE)) (61). Other quinoa seeds come from a carbonaceous dispersion in lens 3z, 700 ± 40 cal BP (UGA-15089, cal. 2σ, 95.4%: 1249-1392 CE) and non-carbon sectors of this same lens dated 760 ± 40 cal BP (UGA-15089, cal. 2σ, 95.4%: 1190-1294 CE).

The locus **Punta de la Peña E (samples #14,15: PPE-w, PPE-d)** corresponds to an archaeological assemblage interpreted as an intentional deposit due to propitiatory practices associated with fertility. It is composed of a ceramic vessel containing a folded textile fragment, covered with a filling of clay sediment of intense reddish color and a high volume of plant material. From this sedimentary filling come numerous intact quinoa seeds dated *ca* 796 ± 24 cal BP (AA-105653, cal. 2σ, 95.4%: 1206-1275 CE). The location of the deposit corresponds to a horizontally raised platform on top of the ignimbrite cliff of Punta de la Peña, at the foot of which there are numerous residential and productive spaces covering a wide temporal sequence.

East of the Puna, **Cueva de Los Corrales 1 site (sample #16: TC2)** is a cave located at 2966 masl in an area of mesothermal valleys, on the west bank of the lower Los Corrales river (El Infiernillo, Tucumán). It comprises a stratigraphic sequence 30 cm thick made of two layers of anthropic origin, separated into three extractions in each case: Layer 1 (1st, 2nd and 3rd extractions) and Layer 2 (1st, 2nd and 3rd extractions). The excellent natural conditions of preservation allowed the recovery of a wide variety of archaeological remains of both inorganic and organic origin. Cueva de Los Corrales 1 has been defined as a multiple activity site with emphasis on processing, consumption and disposal of animal and plant food resources (62). Quinoa seeds analyzed in this paper come from Layer 2 (1st and 2nd extractions) dated *ca* 1270 ± 30 cal BP (UGA-22266, cal. 2σ, 95.4%: 663-859 CE).

The Alero 1 in the **Punta de la Peña 9.I** site **(samples #17, 18: AP9-w, AP9-d)** is located on the edge of the plateau that defines the Sector I of the site, close to a number of stone-walled structures corresponding to agro-pastoralist occupations between *ca* 1500 and 1100 BP. The rocky repair consists of large blocks detached from the ignimbrite cliff of Punta de la Peña. It is apparently collapsed and its walls and top are sooted. The context of interest for this work (Layer 2) lies below a sandy layer (Layer 1), possibly of eolian origin. Layer 2 is composed of a sandy matrix with a wide variety of plant remains recovered in high concentration, together with faunal remains, cordage and scarce lithic and ceramic materials. Macrobotanical remains of *Chenopodium* include dried, non-charred quinoa seeds and fragments of stems and distal ends of the panicles. Quinoa seeds were dated *ca* 1364 ± 20 cal BP (AA-107154, cal. 2σ, 95.4%: 655-766 CE). There are also charred quinoa seeds that constitute waste material from post-harvesting and processing activities, probably for culinary purposes.

**Cueva Salamanca 1 (sample #19: ACS)** is a large cave on the northern margin of the Las Pitas River at an elevation of 3665 masl. The cave is 11 m wide, 8 m deep and 7 m tall (77 m^2^). A total of 30 m^2^ of the site has been excavated so far. Three stone structures are found beside the back wall. The stratigraphy of the cave covers at least 5000 years, and the sediments are the result of natural processes–mostly eolian–and human activity. A series of ten living surfaces that contain hearths, tools, lithic debitage, vegetal remains and grass features have been excavated. Of interest for this paper is the upper stratum, which overlays a lens of volcanic ash. A radiocarbon date *ca* 4500 BP immediately below the volcanic ash gives a *terminus post quem* for the volcanic episode and thus the upper stratum (60). This stratum–level 1(2<u>^a^</u>)–included two hearths, abundant lanceolate nonstemmed obsidian points, a stemmed point of the Punta de la Peña C type (70), grinding stones, quinoa seeds dated *ca* 1796 ± 93 cal BP (AA-107153, cal. 2σ, 95.4%: 231-358 CE), quinoa stems dated *ca* 1742 ± 22 cal BP (AA-107155, cal. 2σ, 95.4%: 250-409 CE), and a spherical pit-like feature that cut through the volcanic ash lens. Above is another stratigraphic unit–level 1(1<u>^a^</u>)–that consists of a loose sandy surface that included ceramic sherds, a small lanceolate and nonstemmed projectile point attributed to the Peñas Chicas E morphological type (70) and three sub-circular stone features that lacked anthropogenic content.

**Table SI-1.** Succinct chronology of climatic and social changes in the Southern dry Andes.

| Years BP | Years CE | Climate | Society | Quinoa samples |
| --- | --- | --- | --- | --- |
| 127 to the present | 1890 to the present | global warming<br>dry phase | continued rural emigration in a context of accelerated technological change, industrialization, urbanization | #1-12 |
| 157 to 127 | 1860 to 1890 | extreme drought | rural depopulation due to mortality and emigration |  |
| 370 to 157 | 17th to 19th centuries | dry phase | efficient crop-pasture systems sustaining regional economy, abandonment of intensified crop fields in Southern highlands due to an emphasis in animal husbandry |  |
| 482 | 1535 | dry phase | beginning of Spanish Conquest facing a century of local rebellion |  |
| 567 to 482 | 1450 to 1535 | dry phase | Inka colonization, continued agricultural intensification |  |
| 1100 to 567 | 850 to 1450 | dry phase | localized agricultural intensification (irrigation, terracing), corporate societies | #13, 14, 15 |
| 1500 to 1100 | 450 to 850 | dry phase | agropastoralist societies | #16, 17, 18 |
| 5000 to 1500 | -3000 to 450 | humid phase | pastoralism, early plant domesticates | #19 |
| 8000 to 5000 | -6000 to -3000 | dry phase | hunter-gatherers |  |
| 12000 to 8000 | -10000 to -6000 | humid phase | hunter-gatherers |  |

### SI-2. SEED SAMPLE DESCRIPTION

**Table SI-2.** Seed sample description.

SI-2. SEED SAMPLE DESCRIPTION **Table SI-2.** Seed sample description.
| N° | Code | Site, Department, Province<sup>a</sup> | Longitude,<br>latitude<sup>b</sup> | Altitude<br>m.a.s.l | Seed count<br>/color<sup>c</sup> | Age<br>cal BP<sup>d</sup> |
| --- | --- | --- | --- | --- | --- | --- |
| 1 | CHEN 458 | Morro de Pucará, Sta Victoria, Salta | -64.97, -22.18 | 2645 | 8/w | modern |
| 2 | CHEN 461 | Poscaya, Sta Victoria, Salta | -65.08, -22.45 | 3208 | 8/w | modern |
| 3 | CHEN 466 | S. José del Aguilar, Sta Victoria, Salta | -65.17, -22.34 | 3960 | 8/w | modern |
| 4 | CHEN 446 | Humahuaca, Jujuy | -65.18, -23.08 | 3823 | 7/w | modern |
| 5 | CHEN 275 | 1485 Coctaca, Humahuaca, Jujuy | -65.28, -23.15 | 3215 | 8/w | modern |
| 6 | CHEN 414 | La Poma, Salta | -66.20, -24.72 | 3016 | 3/w | modern |
| 7 | CHEN 272 | 1482 La Poma, Jujuy | -65.82, -23.85 | 3480 | 8/d | modern |
| 8 | EST | Las Estancias, Andalgalá, CTM | -66.03, -27.58 | 1650 | 8/w | modern |
| 9 | CHEN 427 | Puesto Sey, Susques, Jujuy | -66.48, -23.95 | 4012 | 8/w | modern |
| 10 | BL | Barranca Larga, Belén, CTM | -66.74, -26.98 | 2400 | 8/w | modern |
| 11 | CHEN 420 | Antofallita, Los Andes, Salta | -67.52, -25.25 | 3498 | 5/w | modern |
| 12 | ANT | Antofagasta de la Sierra, CTM | -67.42, -26.05 | 3590 | 8/w | modern |
| 13 | AP4 | Punta de la Peña 4, ADLS, CTM | -67.33, -26.02 | 3590 | 10/w | 690 ± 50 |
| 14 | PPE-w | Punta de la Peña E, ADLS, CTM | -67.33, -26.02 | 3590 | 19/w | 796 ± 24 |
| 15 | PPE-d | Punta de la Peña E, ADLS, CTM | -67.33, -26.02 | 3590 | 10/d | 796 ± 24 |
| 16 | TC2 | Cueva Corrales 1, Tafí del Valle, TUC | -65.80, -26.73 | 2966 | 2/w | 1270 ± 30 |
| 17 | AP9-w | Punta de la Peña 9, ADLS, CTM | -67.33, -26.02 | 3590 | 25/w | 1364 ± 20 |
| 18 | AP9-d | Punta de la Peña 9, ADLS, CTM | -67.33, -26.02 | 3590 | 8/d | 1364 ± 20 |
| 19 | ACS | Cueva Salamanca 1, ADLS, CTM | -67.33, -26.01 | 3665 | 2/w | 1796 ± 23 |

<sup>a</sup> ADLS: Antofagasta de la Sierra, CTM: Catamarca, TUC: Tucumán;
<sup>b</sup> longitude and latitude in decimal degree values;
<sup>c</sup> seed count in sample / seed color is dark (d) or white (w);

### SI-3. DNA EXTRACTION, SIMPLE SEQUENCE REPEAT GENOTYPING, ANCIENT DNA QUALITY CONTROL, AND POPULATION GENETIC ANALYSIS

#### Prevention of contamination

Following recommendations to minimize the risk of exogenous DNA contamination and ensure the reliability of the results (71), dissections, extractions and pre-PCR processing of ancient and modern seeds were rigorously separated in time and space. We first processed ancient seeds in a specific laboratory dedicated to ancient DNA under sterile conditions. In the ancient DNA laboratory, we purchased and used new consumables and extraction kits, with the room cleaned and exposed to UV overnight after each DNA extraction cycle, in order to destroy possible traces of DNA between successive extractions. We wore protective clothing and footwear. Once all the extractions and amplifications of archaeological seeds were completed, we then proceeded to the extraction and amplification of modern seeds in a distant laboratory, without any spatial connection with the previous one.

#### Genotyping

We worked on 81 modern and 144 ancient quinoa seeds for DNA extraction according to the below-described procedures. In modern as well as ancient seeds, we dissected seeds under a stereomicroscope (Leica MZ 16, Leica camera DFC 280) to separate the embryo from the central perisperm. We next extracted total DNA from embryos. Quinoa embryos (ca. 1-3 mm length, 1 mm thick) were dissected one seed after the other, using sterile dissection equipment and binoculars. DNA extraction was successful for all the modern seeds, while we recovered well-preserved DNA from only 76 ancient seeds (53%). This level of ancient DNA recovery reflects the high preservation of genetic material in dry environments as pointed out by (72), conditions still improved in the dry Andes by cold temperatures and oxygen scarcity at high altitude (73, 74). Despite these favorable conditions, there appears to be a time limit for the preservation of quinoa seeds in archaeological contexts (75).

#### DNA extraction

Total DNA extraction was obtained using DNeasy Plant Kit (Qiagen, Hilden, Germany) following the DNeasy Tissue Kit Handbook protocol with two 50 μL final elution. We extracted DNA by sets of no more than 12 samples per half-day, with one negative extraction control for each set of extractions.

#### PCR amplification

All quinoa samples were initially genotyped at 25 polymorphic microsatellite loci described by Mason et al. (Mason et al. 2005) and Jarvis et al. (Jarvis et al. 2008). PCR consisted of a final volume of 10 μL containing 0.2 μM of each primer, 2 or 4 μL of DNA solution (depending on the storage quality of the sample), and 5 μL of kit multiplex PCR kit (Qiagen). The amplification parameters were 15 min of 95 °C followed by 30 cycles (40 cycles for ancient quinoa samples) of 94 °C for 60 s, 56 °C for 120 s, and 72 °C for 60 s, with a final extension step for 30 min of 60 °C in a Mastercycler epgradients (Eppendorf). We systematically performed negative controls to check for possible contamination. We independently amplified each individual two times, retaining only congruent results. Amplifications products were separated on an ABI-3100 Automated Sequencer at the SFR SEM platform and analyzed with Genemapper 4.0 (Applied Biosystem) using Genescan-500LIZ size standard (Applied) with two investigators eye checking for allele scoring. Genetic analyses included only reproducible alleles, present in both replicates, and accessions with reliable information for at least 24 of the 25 loci. We discarded locus QAAT100 because of its complex motive and size results out of range). Microsatellite loci, expected size and observed size range, number of alleles per locus and missing data are included in **Tables SI-3.1** and **SI-3.2**.

**Table SI-3.1.** Microsatellite loci, allele expected size, observed size range, allele number, and missing data points in modern (n=81) and ancient (n=76 successfully genotyped out of 144) quinoa seed samples. All loci are derived from (76) except KGA 003 and KGA 20 derived from (77).

| Locus | Expected size (bp) | Observed size (bp) | Allele number | Missing modern data points | Missing ancient data points |
| --- | --- | --- | --- | --- | --- |
| KGA003 | 150 | 139-174 | 12 | 1 | 0 |
| KGA020 | 177 | 152-204 | 22 | 1 | 3 |
| QAAT001 | 182 | 131-225 | 16 | 5 | 9 |
| QAAT011 | 197 | 162-237 | 24 | 5 | 8 |
| QAAT022 | 194 | 138-253 | 31 | 2 | 1 |
| QAAT024 | 198 | 169-229 | 19 | 1 | 1 |
| QAAT026 | 181 | 172-253 | 17 | 20 | 4 |
| QAAT027 | 165 | 147-215 | 20 | 4 | 7 |
| QAAT050 | 199 | 186-242 | 26 | 0 | 1 |
| QAAT062 | 187 | 157-215 | 17 | 7 | 1 |
| QAAT071 | 170 | 130-309 | 37 | 2 | 1 |
| QAAT074 | 186 | 160-218 | 15 | 5 | 12 |
| QAAT078 | 196 | 176-253 | 14 | 4 | 3 |
| QAAT087 | 185 | 165-211 | 14 | 0 | 1 |
| QAAT088 | 151 | 99-173 | 21 | 0 | 46 |
| QAAT097 | 177 | 161-215 | 18 | 1 | 13 |
| QAAT106 | 299 | 288-325 | 11 | 0 | 22 |
| QAAT112 | 199 | 176-227 | 13 | 1 | 4 |
| QATG064 | 177 | 167-182 | 5 | 0 | 4 |
| QCA037 | 188 | 173-194 | 10 | 1 | 22 |
| QCA053 | 189 | 166-200 | 13 | 3 | 0 |
| QCA057 | 163 | 155-191 | 11 | 1 | 6 |
| QGA003 | 150 | 125-197 | 25 | 1 | 0 |
| QGA024 | 195 | 169-229 | 19 | 1 | 1 |
| QAAT100 | 349 | *out of range* | *complex motive* | *discarded* | *discarded* |

**Table SI-3.2.**
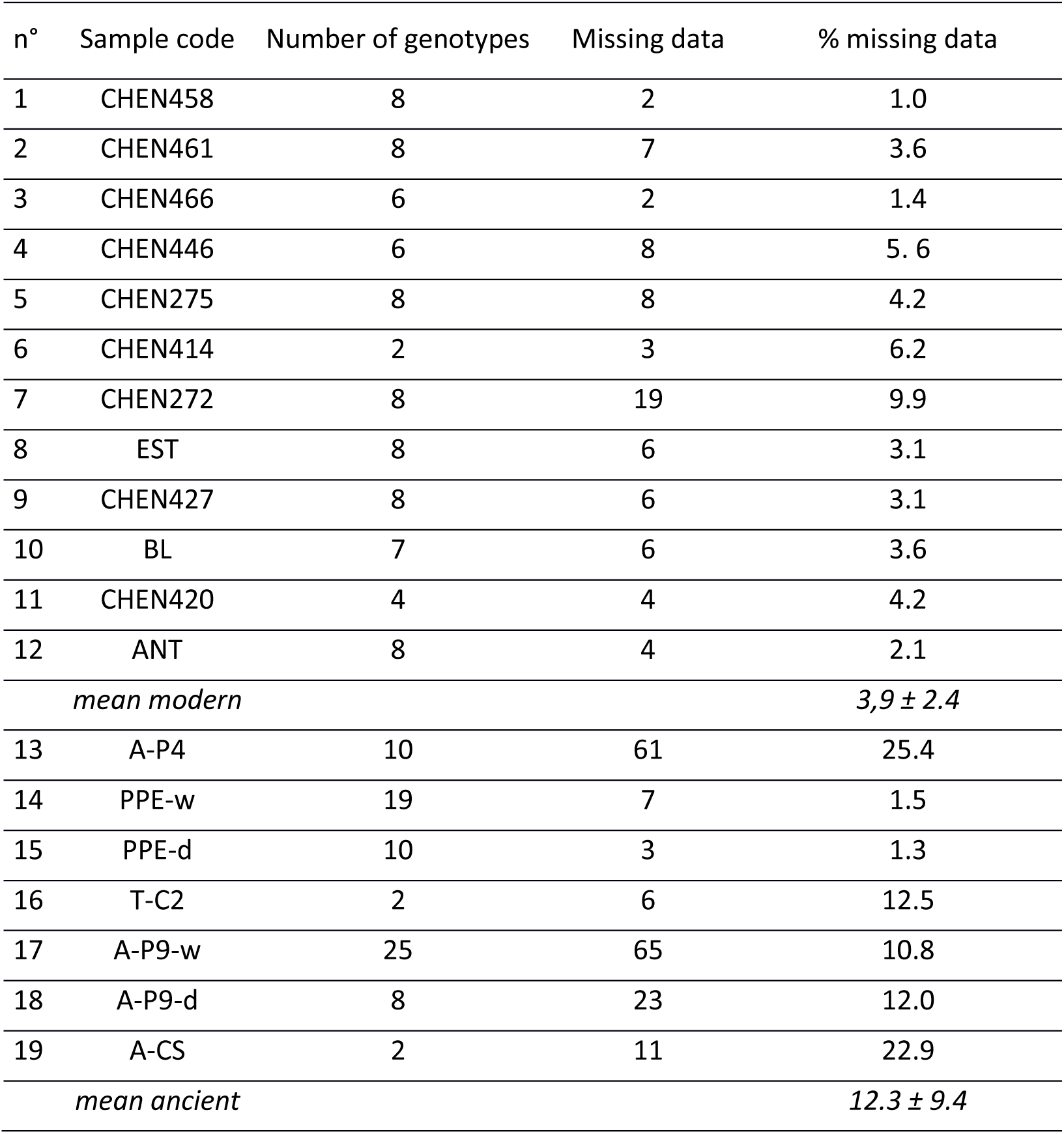
Percentage of missing data per successfully genotyped quinoa seed sample, calculated across 24 microsatellite loci. (#1-12: modern, n=81; #13-19: ancient, n=76).

#### Preliminary analyses of ancient DNA quality

To check for DNA quality in ancient quinoa seeds, alleles from locus QAAT024 and QAAT087 of ancient and modern samples were direct sequenced in forward and reverse sense, using the ABI Prism BigDye Terminator Cycle Sequencing Kit 3.1 in an Applied Biosystems 3500 DNA Sequencer. Reactions containing fragments of the expected size were purified by treatment with Exonuclease I and Shrimp Alkaline Phosphatase. Enzymes were added directly to the PCR product to degrade primers and dephosphorylate dNTPs that were not consumed in the reaction and could interfere with downstream sequencing. Treatment was carried out for 15 minutes at 37 °C, followed by a 15-minute incubation at 80 °C to completely inactivate both enzymes. Base assignment was made with GeneMapper V3.0 software (Applied Biosystems). Phred quality score was settled at 20 to assure 99% of base call accuracy, as a measure of the quality of the identification of the nucleobases generated by automated DNA sequencing. Sequences were aligned using CLUSTALW (78) followed by minor manual modifications. We analyzed amplification products by comparison with the public sequence databases as nucleotide using BLASTN. Following the criteria of Meyers et al. (79), a sequence was classified as a known element when retrieved with a BLAST E-value of less than 10^−5^.

Sequence analysis revealed amplification products corresponding to what was expected for both microsatellite loci. After BLAST search, we found modern sequences to have 100% homology with *Chenopodium quinoa* clones of microsatellite sequences with E-values of virtually zero. At locus QAAT024, we selected 6 ancient and 4 modern genotypes, 2 ancient samples failed to give positive results. Finally, we obtained 8 consensus sequences, from 4 modern and 4 ancient genotypes (**Fig SI-3.1**). In the microsatellite region, a six-base pair InDel was present between these modern genotypes, according to the size of the expected fragment. The ancient sequences revealed variations at a total of seven nucleotide positions, representing 97% of sequence similarity with modern sequences. Variation at nucleotide position 70 was on only one single base in one ancient genotype. The other 6 mutations discriminated between modern and ancient sample groups, as well as among ancient genotypes (nucleotide position 229), which confirms the absence of contaminating sequences.

**Fig SI-3.1.**
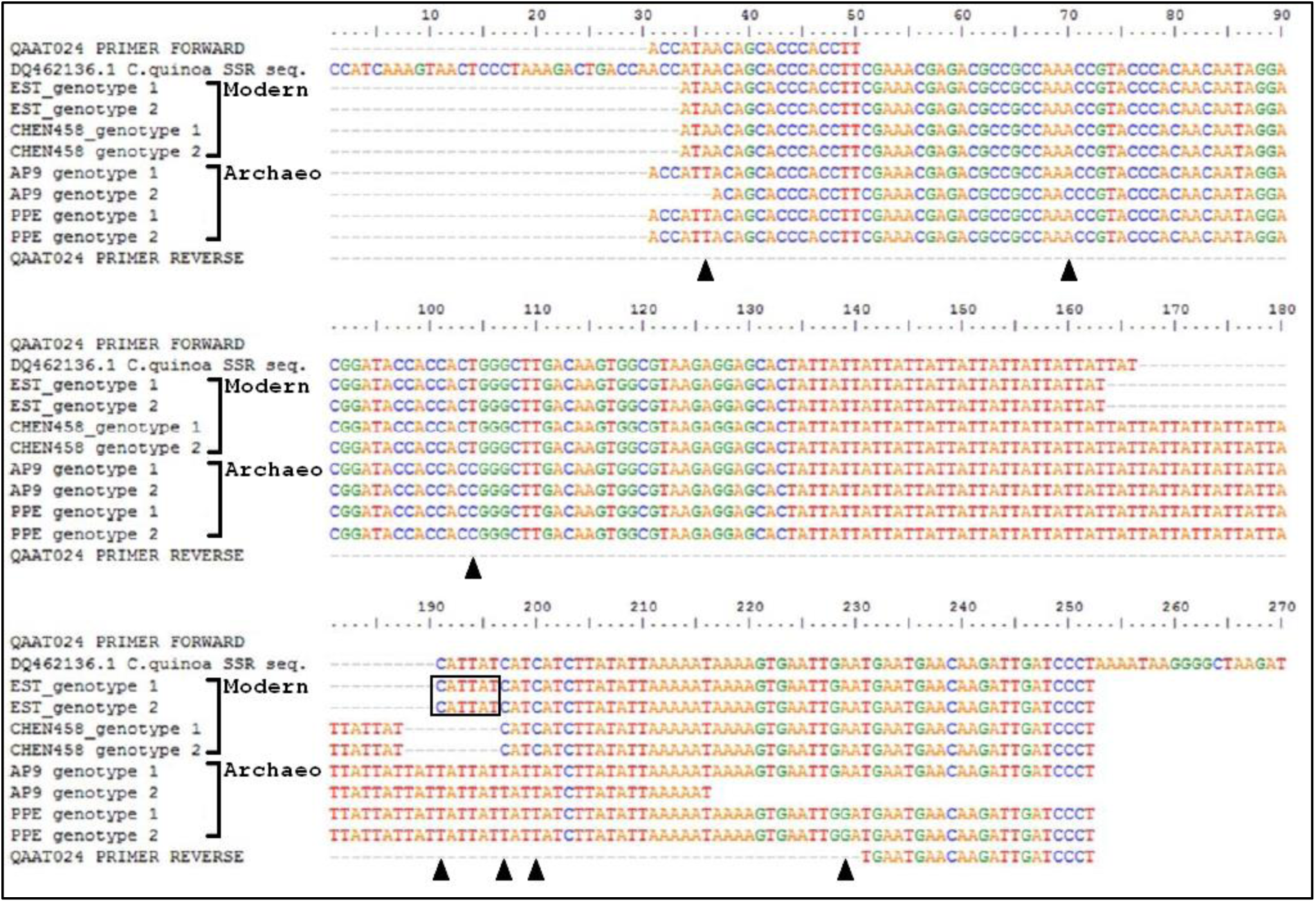
Sequence alignment from QAAT024 locus alleles of 4 modern (EST and CHEN458 samples) and 4 ancient (AP9 and PPE samples) quinoa genotypes. Primer and NCBI accession sequences are included. Arrows show sequence variations, box shows the 6-base pair InDel.

At locus QAAT087, we obtained 4 consensus sequences, from 3 ancient and 1 modern genotype (the remaining samples failed to give positive results) (**Fig SI-3.2**). Sequences from locus QAAT087 have a lower quality than locus QAAT024, not only in archeological samples but also in modern ones. Alignment analyses revealed sequence variations at a total of 7 nucleotide positions, representing 97% of sequence similarity with modern sequences.

**Fig SI-3.2.**
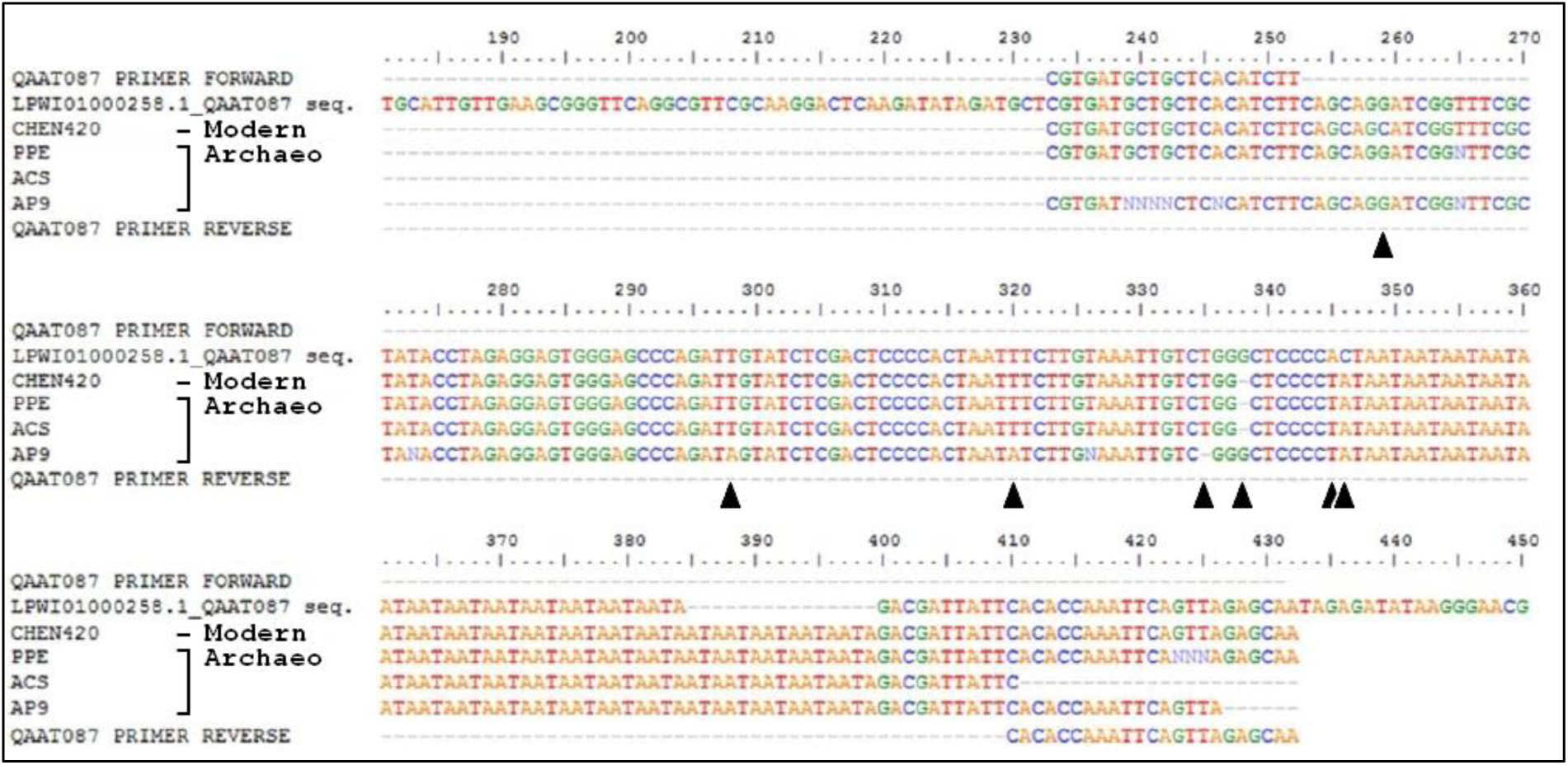
Sequence alignment from QAAT087 locus alleles of one modern (CHEN 420) and three ancient (PPE, ACS, AP9) genotypes. Primer and NCBI accession sequences are included. Arrows show sequence variations.

Allele identities were corroborated by sequencing and CLUSTAL alignment and BLAST algorithms. Amplifications were successful for most of the samples, indicating an adequate quality of both ancient and modern DNA. Taking into account that all caution was taken to avoid contamination, our results agree with O’Donogue (72), who suggests that the microenvironment within desiccated seeds is conducive to enhanced preservation of lipids, nucleic acids and other biomolecules.

#### Genetic diversity indexes

We measured the genetic diversity of each sample using the allelic richness *N_all-rar_* (80) and the expected heterozygosity *H_e_*. In predominantly selfing populations, we expect a strong deviation compared to Hardy-Weinberg expectations. We estimated the inbreeding fixation coefficient *F*_IS_ for each sample and *F*_ST_ for each pair of samples according to Weir & Cockerham (81). *F*_IS_ and *F*_ST_ measure genetic differentiation within and among populations respectively; both coefficients range from zero (no differentiation) to one (complete differentiation). Analyses were performed in R using the packages *adegenet* and *hierfstat* (63, 64) and the program ADZE for rarefaction analyses (66).

We expect quinoa populations to be highly selfing and therefore to display a limited number of repeated multilocus genotypes (thereafter MLG). We used the package *poppr* to count the number of MLGs present in each population (*nbMLG*) (65). Populations with no more than two individuals sampled were removed from this analysis. We then used the rarefaction method (ADZE) to estimate the expected MLG richness (*eMLG*) corrected for the differences in sample size. Finally, within each population, the composition in MLGs can either be balanced, or highly biased with one predominant MLG. We measured this using the Simpson diversity index (λ) that is equivalent to a multilocus expected heterozygosity.

#### Selfing rate estimation

Two independent estimates of selfing rates were calculated: either directly as *s*(*F_Is_*)*=2 F*_Is_/(1+*F*_Is_) (82), or as *s*(*LnL*) using the program RMES (robust multilocus estimate of selfing), based on the distribution of multilocus heterozygosity, which allowed calculating a confidence interval at P=95% (83). We used the maximum likelihood estimation with a precision of 0.00001, a maximum number of generations of selfing set to 10 and 100000 iterations. In some populations, the high degree of homozygosity (#1,3,6) or the low sample size (samples #10,11,16,19) prevented the estimation of *s*(*LnL*). Results of allelic diversity, heterozygosity and selfing rates in the 19 studied quinoa samples are shown in **Table SI-3.3**.

**Table SI-3.3.** Diversity and selfing rates in the 19 studied quinoa seed samples. (samples #1-12: modern; #13-19: ancient).

| *n*<sup>o</sup> | *sample* | *N* | *N<sub>all-rar</sub>* | *H<sub>e</sub>* | *nbMLG* | *eMLG* | $\lambda$ | *F<sub>IS</sub>* | *s(F<sub>IS</sub>)* | *s(LnL)* | *CI (95%)* |
| --- | --- | --- | --- | --- | --- | --- | --- | --- | --- | --- | --- |
| 1 | CHEN458 | 8 | 1.00 | 0.02 | 3 | 2 | 0.406 | 1.00 | 1.00 | - | - |
| 2 | CHEN461 | 8 | 1.94 | 0.39 | 8 | 4 | 0.875 | 0.50 | 0.67 | 0.52 | 0.31 - 0.71 |
| 3 | CHEN466 | 6 | 1.10 | 0.07 | 5 | 3.6 | 0.778 | 0.80 | 0.89 | - | - |
| 4 | CHEN446 | 6 | 1.39 | 0.18 | 6 | 4 | 0.833 | 0.60 | 0.75 | 0.89 | 0.82 - 0.94 |
| 5 | CHEN275 | 8 | 2.39 | 0.56 | 8 | 4 | 0.875 | 0.15 | 0.26 | 0.21 | 0.12 - 0.37 |
| 6 | CHEN414 | 2 | - | 0.05 | - | - | - | 1.00 | 1.00 | - | - |
| 7 | CHEN272 | 8 | 3.05 | 0.70 | 8 | 4 | 0.875 | 0.02 | 0.04 | 0.12 | 0.03 - 0.24 |
| 8 | EST | 8 | 2.51 | 0.61 | 8 | 4 | 0.875 | 0.81 | 0.90 | 0.42 | 0.01 - 0.73 |
| 9 | CHEN427 | 8 | 2.76 | 0.67 | 8 | 4 | 0.875 | 0.58 | 0.73 | 0.72 | 0.68 - 0.79 |
| 10 | BL | 7 | 1.44 | 0.17 | 7 | 4 | 0.857 | 0.51 | 0.68 | - | - |
| 11 | CHEN420 | 4 | 1.36 | 0.17 | 4 | 4 | 0.750 | 0.82 | 0.90 | - | - |
| 12 | ANT | 8 | 1.81 | 0.36 | 8 | 4 | 0.875 | 0.28 | 0.44 | 0.69 | 0.65 - 0.77 |
| 13 | AP4 | 10 | 2.94 | 0.71 | 10 | 4 | 0.900 | 0.66 | 0.80 | 0.75 | 0.68 - 0.82 |
| 14 | PPE-w | 19 | 2.69 | 0.61 | 19 | 4 | 0.947 | 0.66 | 0.80 | 0.76 | 0.73 - 0.80 |
| 15 | PPE-d | 10 | 2.77 | 0.66 | 10 | 4 | 0.900 | 0.54 | 0.70 | 0.78 | 0.75 - 0.82 |
| 16 | TC2 | 2 | - | 0.95 | - | - | - | 0.97 | 0.98 | - | - |
| 17 | AP9-w | 25 | 3.05 | 0.72 | 25 | 4 | 0.960 | 0.48 | 0.65 | 0.54 | 0.45 - 0.61 |
| 18 | AP9-d | 8 | 3.11 | 0.74 | 8 | 4 | 0.875 | 0.35 | 0.52 | 0.59 | 0.55 - 0.69 |
| 19 | ACS | 2 | - | 0.70 | - | - | - | 0.36 | 0.53 | - | - |

*N*: number of individuals; *N<sub>all-rar</sub>*: allelic richness estimated using ADZE; *H<sub>e</sub>*: expected heterozygosity; *nbMLG*: number of multilocus genotypes observed; *eMLG*: MLG richness estimated by a rarefaction method; $\lambda$ : Simpson diversity index; *F<sub>IS</sub>*: inbreeding fixation coefficient; *s(F<sub>IS</sub>)*: selfing rate estimated from the *F<sub>IS</sub>*; *s(LnL)*: selfing rate estimated by RMES ; *CI (95%)*: confidence interval of *s(LnL)* at *P*=95%.

**Table SI-3.4.**
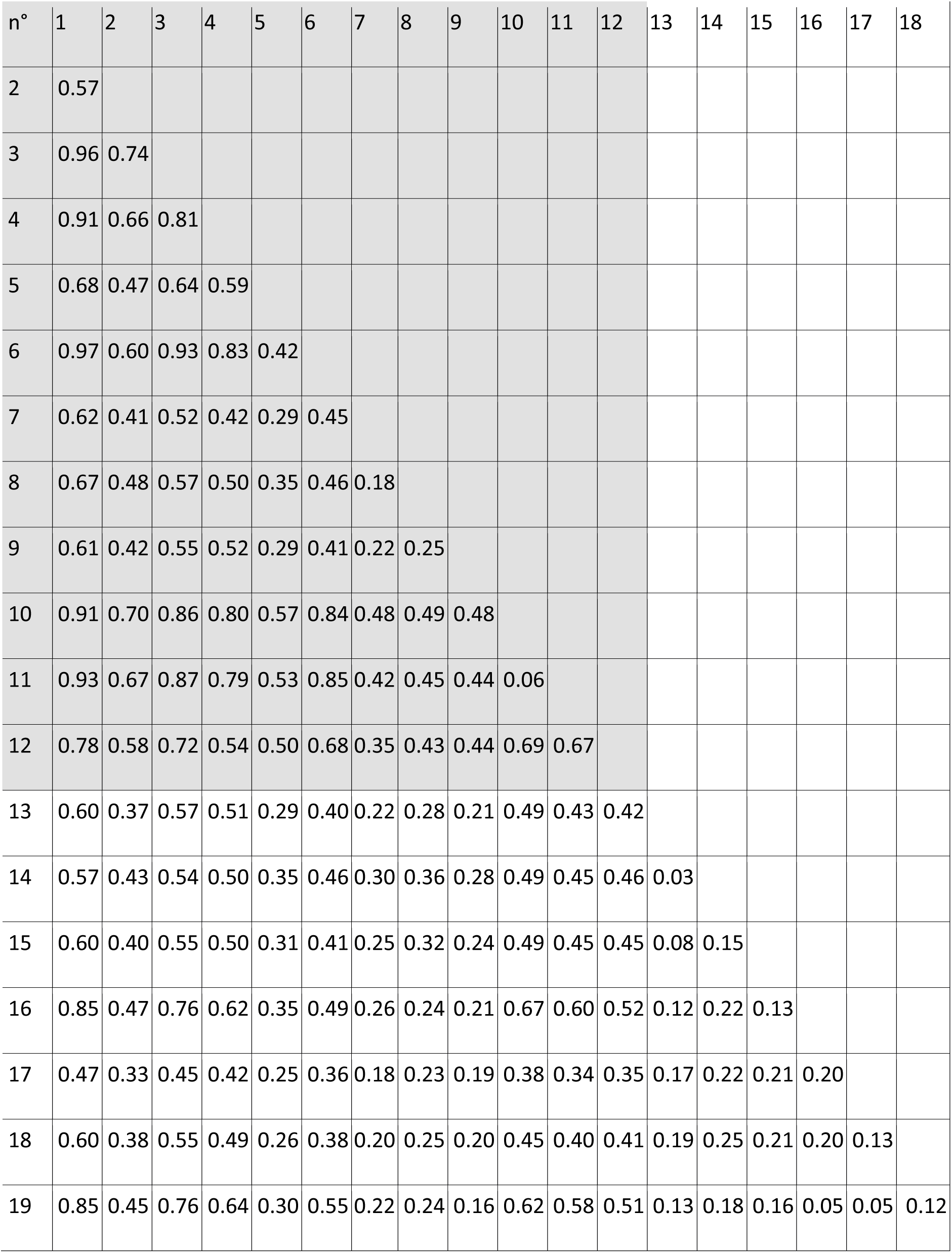
Pairwise *F*_ST_ values between samples. (samples #1-12: modern; #13-19: ancient).

| n° | 1 | 2 | 3 | 4 | 5 | 6 | 7 | 8 | 9 | 10 | 11 | 12 | 13 | 14 | 15 | 16 | 17 | 18 |
| --- | --- | --- | --- | --- | --- | --- | --- | --- | --- | --- | --- | --- | --- | --- | --- | --- | --- | --- |
| 2 | 0.57 |  |  |  |  |  |  |  |  |  |  |  |  |  |  |  |  |  |
| 3 | 0.96 | 0.74 |  |  |  |  |  |  |  |  |  |  |  |  |  |  |  |  |
| 4 | 0.91 | 0.66 | 0.81 |  |  |  |  |  |  |  |  |  |  |  |  |  |  |  |
| 5 | 0.68 | 0.47 | 0.64 | 0.59 |  |  |  |  |  |  |  |  |  |  |  |  |  |  |
| 6 | 0.97 | 0.60 | 0.93 | 0.83 | 0.42 |  |  |  |  |  |  |  |  |  |  |  |  |  |
| 7 | 0.62 | 0.41 | 0.52 | 0.42 | 0.29 | 0.45 |  |  |  |  |  |  |  |  |  |  |  |  |
| 8 | 0.67 | 0.48 | 0.57 | 0.50 | 0.35 | 0.46 | 0.18 |  |  |  |  |  |  |  |  |  |  |  |
| 9 | 0.61 | 0.42 | 0.55 | 0.52 | 0.29 | 0.41 | 0.22 | 0.25 |  |  |  |  |  |  |  |  |  |  |
| 10 | 0.91 | 0.70 | 0.86 | 0.80 | 0.57 | 0.84 | 0.48 | 0.49 | 0.48 |  |  |  |  |  |  |  |  |  |
| 11 | 0.93 | 0.67 | 0.87 | 0.79 | 0.53 | 0.85 | 0.42 | 0.45 | 0.44 | 0.06 |  |  |  |  |  |  |  |  |
| 12 | 0.78 | 0.58 | 0.72 | 0.54 | 0.50 | 0.68 | 0.35 | 0.43 | 0.44 | 0.69 | 0.67 |  |  |  |  |  |  |  |
| 13 | 0.60 | 0.37 | 0.57 | 0.51 | 0.29 | 0.40 | 0.22 | 0.28 | 0.21 | 0.49 | 0.43 | 0.42 |  |  |  |  |  |  |
| 14 | 0.57 | 0.43 | 0.54 | 0.50 | 0.35 | 0.46 | 0.30 | 0.36 | 0.28 | 0.49 | 0.45 | 0.46 | 0.03 |  |  |  |  |  |
| 15 | 0.60 | 0.40 | 0.55 | 0.50 | 0.31 | 0.41 | 0.25 | 0.32 | 0.24 | 0.49 | 0.45 | 0.45 | 0.08 | 0.15 |  |  |  |  |
| 16 | 0.85 | 0.47 | 0.76 | 0.62 | 0.35 | 0.49 | 0.26 | 0.24 | 0.21 | 0.67 | 0.60 | 0.52 | 0.12 | 0.22 | 0.13 |  |  |  |
| 17 | 0.47 | 0.33 | 0.45 | 0.42 | 0.25 | 0.36 | 0.18 | 0.23 | 0.19 | 0.38 | 0.34 | 0.35 | 0.17 | 0.22 | 0.21 | 0.20 |  |  |
| 18 | 0.60 | 0.38 | 0.55 | 0.49 | 0.26 | 0.38 | 0.20 | 0.25 | 0.20 | 0.45 | 0.40 | 0.41 | 0.19 | 0.25 | 0.21 | 0.20 | 0.13 |  |
| 19 | 0.85 | 0.45 | 0.76 | 0.64 | 0.30 | 0.55 | 0.22 | 0.24 | 0.16 | 0.62 | 0.58 | 0.51 | 0.13 | 0.18 | 0.16 | 0.05 | 0.05 | 0.12 |

#### Genetic structure

We investigated the genetic structure among the samples with a multivariate approach, the Discriminant analysis of Principal Components (DAPC) implemented in the package *adegenet* in R environment (84) (**Fig 1B** in main text). To avoid over-fitting the model, we used the cross validation method to choose the optimum number of principal components to include in the model. We retained 15 principal components and 4 discriminant factors in the final model, which explained 57% of the sample variability. To find the number of genetic groups better fitting our sample, we used the k-means algorithm for a number of groups k=1−25. We ran 10^9^ iterations with 2,000 starting points. The Bayesian Information Criterion (BIC) minimized at k=12 for the whole sample analysis (**Figs. SI-3.3, SI-3.4**).

**Fig SI-3.3.**
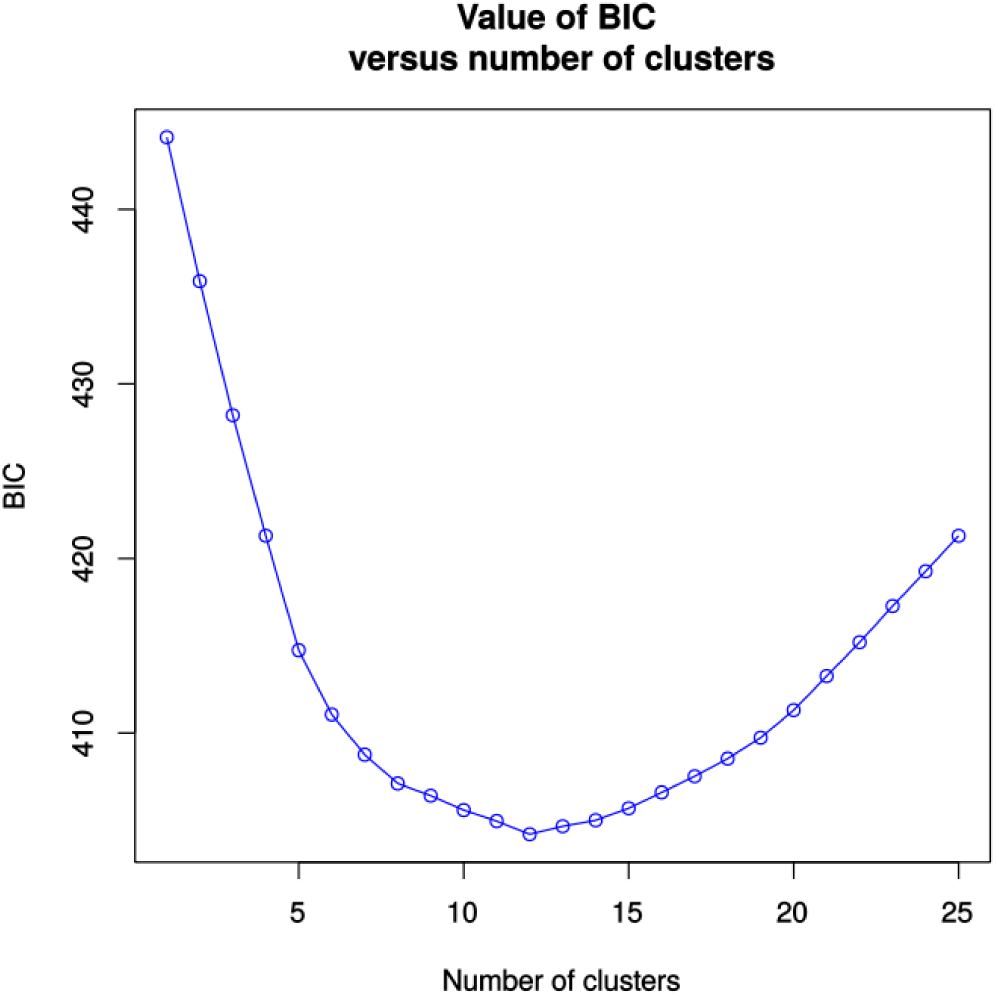
Inference of number of clusters in DAPC. The Bayesian Information Criterion (BIC) is minimum at k=12, suggesting an optimal separation of samples in 12 groups.

**Fig SI-3.4.**
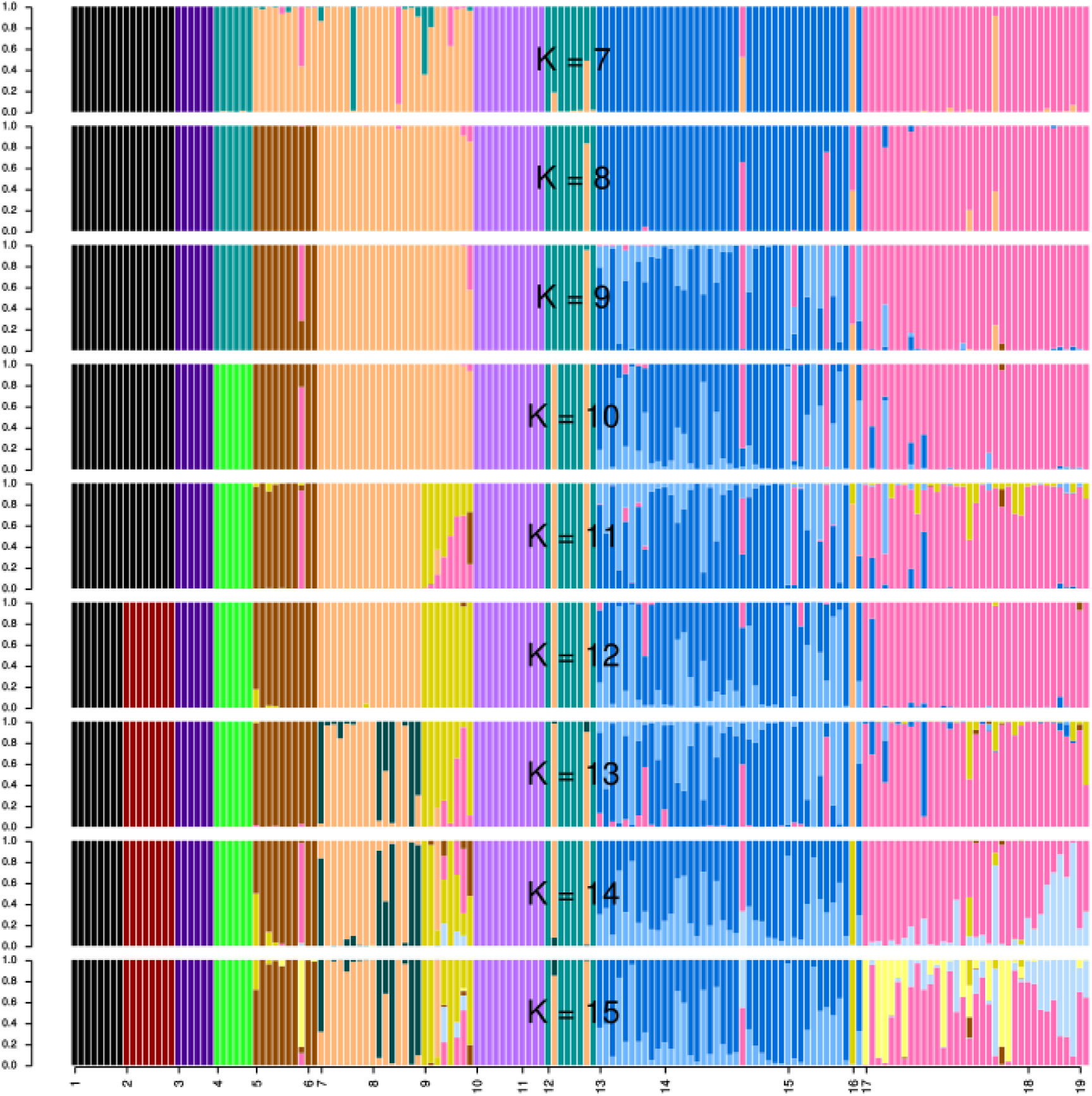
Individual assignment probability to each group from Discriminant Analysis of Principal Components (DAPC). Horizontal axis shows the population codes as in **Table SI-2**. Vertical axis shows the assignment probabilities for values of k from 7 to 15. Colors for k=12 are the same as in **Fig 1B** in main text.

We also used a Bayesian clustering approach implemented in the program STRUCTURE v2.3.4 using the correlated-frequencies model without admixture (67) (**Fig SI-3.5**). We assessed the structure of the sample for a number of clusters k=7−15. For each value of k, we ran 10 repetitions of 2×10^6^ iterations of the MCMC algorithm after discarding 5×10^5^ iterations as burn-in. Finally, we performed a Principal Component Analysis with the *adegenet* package (64) (**Fig SI-3.6**).

**Fig SI-3.5.**
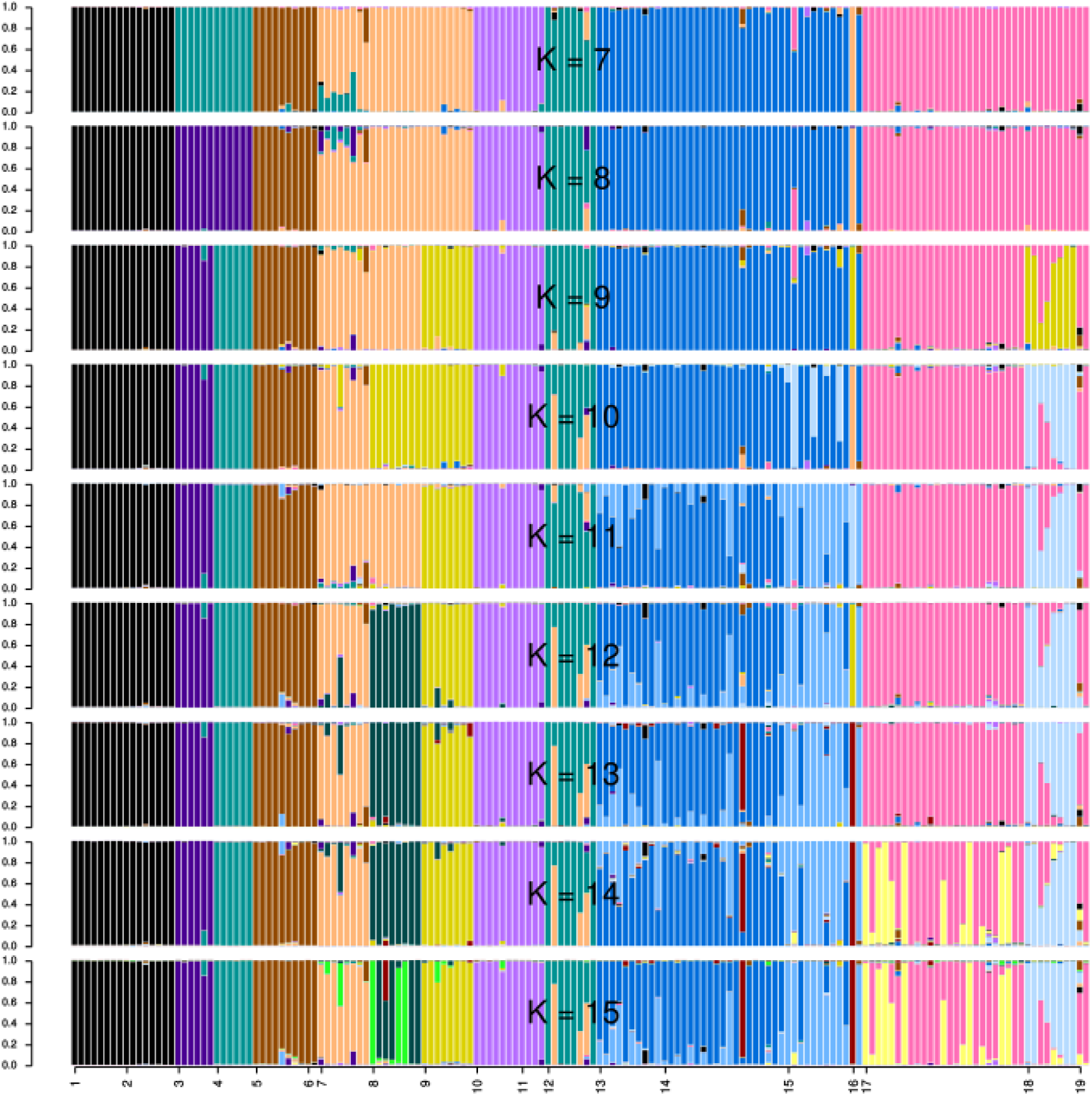
Individual assignment probability to each group from STRUCTURE. Horizontal axis shows the population codes as in **Table SI-2**. Vertical axis shows the assignment probabilities for values of k from 7 to 15. For each value of k only the run with highest likelihood is represented. Colors for k=12 are the same as in **Fig SI-3.6** showing the results from PCA.

**Fig SI-3.6.**
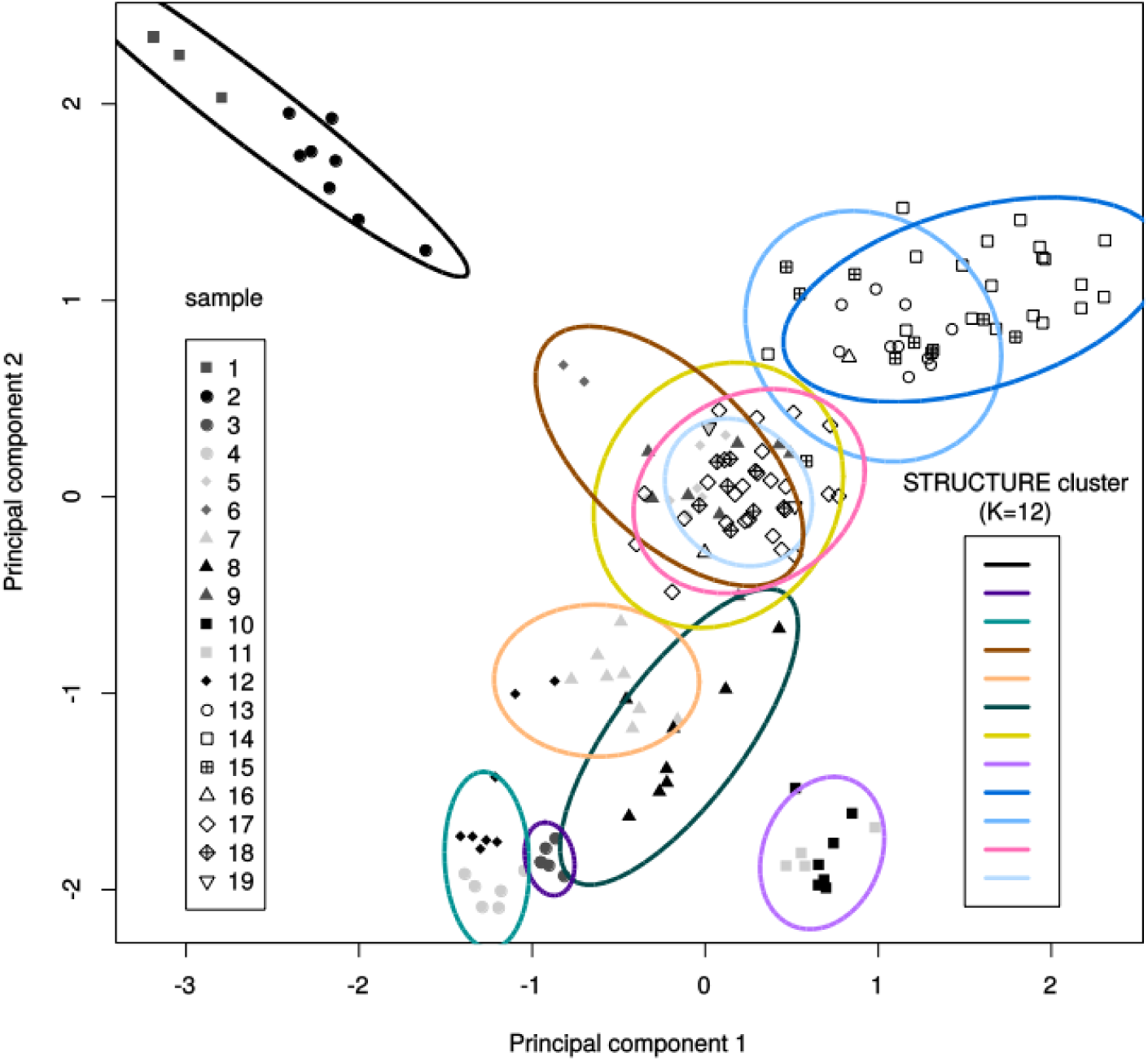
First two principal components for the Principal Component Analysis (PCA). Colored ellipses show the genetic groups identified with STRUCTURE for k=12 (see **Fig SI-3.5**).

#### Coalescent-based analyses

An approximate Bayesian computation (ABC) approach using random forests (68, 69) was used to characterize the local demographic history of quinoa found around Antofagasta de la Sierra. The coalescence-based modeling of genetic relationships between modern and ancient samples from Antofagasta examined six scenarios (**Fig 2**): (i) direct chronological filiation between all samples, mixing cultivated and wild forms (**Fig 2A**), ii) replacement of all ancient quinoas by modern quinoas (**Fig 2B**), iii) filiation between modern quinoas and intermediate ancient quinoas, both replacing the oldest quinoas (**Fig 2C**), iv) successive replacement of the three groups of quinoas: modern, intermediate, ancient (**Fig 2D**), v) same scenario as previously but differentiating between cultivated and wild forms (**Fig 2E**), vi) same scenario as previously but with admixture between cultivated and wild forms (**Fig 2F**).

Coalescent simulations and calculation of summary statistics were performed with DIYABC (85). Reference tables were exported and ABC model choice and parameter estimation was performed with the random forest approach implemented in the R package *abcrf* (86). The scripts in R employed are available at the Zenodo open access repository (https://zenodo.org/). All single-sample and two-sample summary statistics available at DIYABC and admixture summary statistics (from a reduced number of relevant sample trios) were used to grow random forests for ABC. **Table SI-3.5**. presents prior and posterior probability distributions for parameters of the models. For each scenario 60 000 simulations were performed. Random forests of 800 trees were grown for model choice. For the best model, 140 000 additional simulation were run and random forests of 1000 trees were grown for parameter estimation.

**Table SI-3.5.** Prior and posterior probability distribution of model parameters for the approximate Bayesian computation analysis.

| Parameter | Prior distribution | conditions | Posterior (median and 95%HPD)* |
| --- | --- | --- | --- |
| Age sample #12** | Constant=-56 |  |  |
| Age sample #13 ($t_1$)** | Normal (m=690, $\sigma$=50) | | Indistinguishable from prior |
| Age samples #14–15 ($t_2$)** | Normal (m=796, $\sigma$=24) | $t_2 > t_1$ | Indistinguishable from prior |
| Age samples #17–18 ($t_3$)** | Normal (m=1364, $\sigma$=20) | $t_3 > t_2$ | Indistinguishable from prior |
| Age sample #19 ($t_4$)** | Normal (m=1796, $\sigma$=23) | $t_4 > t_3$ | Indistinguishable from prior |
| Admixture proportion from wild genetic pool for sample #15 ($\alpha_1$) | Uniform (min=0, max=1) | | 0.35 (0.18–0.69) |
| Admixture proportion from wild genetic pool for sample #18 ($\alpha_2$) | | | 0.48 (0.30–0.82) |
| Time of divergence ($td_1$), between clusters | Uniform (min=2000, max=7000) | $td_2 > td_1$ | Indistinguishable from prior |
| Time of domestication ($td_2$), domesticated and wild | Uniform (min=5000, max=7000) | | Indistinguishable from prior |
| Effective population size ($Na_0$), ‘Green’ cluster (sample #12) | Log-Uniform (min=2, max=$10^6$) | | 185 (37–1231) |
| Effective population size ($Nb_0$), ‘Blue’ cluster (samples #13–15) between $t_1$ and $t_2$ | | | 866 (133–64026) |
| Effective population size ($Nb_1$), ‘Blue’ cluster (samples #13–15) between $t_2$ and $td_1$ | | | 857 (195–2981) |
| Effective population size ($Nc_0$), ‘Red’ cluster (samples #17–19) between $t_3$ and $t_4$ | | | 1469 (535–7587) |
| Effective population size ($Nc_1$), ‘Red’ cluster (samples #17–19) between $t_4$ and $td_1$ | | | 1035 (130–116845) |
| Effective population size ($N_w$), wild population | | | 30 (2–3094) |
| Other effective population size parameters (e.g. ancestral populations) |  |  | Indistinguishable from prior |
| Mutation rate ($\mu$) | Log-Uniform (min=$10^{-5}$, max=$10^{-2}$) | | $1.10 \times 10^{-3}$ ($8.18 \times 10^{-5}$–$5.59 \times 10^{-3}$) |
| Geometric distribution parameter for GSM ($P_{GSM}$) | Uniform (min=0, max=1) | | 0.60 (0.09–0.98) |
| Mutation rate SNI ($\mu_{SNI}$) | Log-Uniform (min=$10^{-9}$, max=$10^{-5}$) | | Indistinguishable from prior |

\* Only reported posterior probability estimates that differ conspicuously from prior probability distributions.
\*\* Time measured in years before present, ‘present’ being set to calendar year 1950 to follow scale of radiocarbon dating.

## References

1. Binford MW, Kolata AL, Brenner M, Janusek JW, Seddon MT, Abbott M, et al. Climate variation and the rise and fall of an Andean civilization. Quatern Int. 1997, 47: 235–48.

2. Contreras DA. Landscape and Environment: Insights from the Prehispanic Central Andes. J Archaeol Res. 2010, 18: 241–88.

3. Ortloff CR, Kolata AL. Climate and collapse - agroecological perspectives on the decline of the Tiwanaku state. J Archaeol Sci. 1993, 20: 195–221.

4. Craig N. Cultural dynamics, climate, and landscape in the South-Central Andes during the mid-late Holocene: a consideration of two socio-natural perspectives. Chungara. 2011, 43: 367–91.

5. Winkel T, Bommel P, Chevarría-Lazo M, Cortes G, Del Castillo C, Gasselin P, et al. Panarchy of an indigenous agroecosystem in the globalized market: The quinoa production in the Bolivian Altiplano. Glob Environ Change. 2016, 39: 195–204.

6. Zimmerer KS. The compatibility of agricultural intensification in a global hotspot of smallholder agrobiodiversity (Bolivia). Proc Natl Acad Sci USA. 2013, 110: 2769–74.

7. Pearsall DM. Plant domestication and the shift to agriculture in the Andes. In: Silverman H, Isbell WH, editors. The Handbook of South American Archaeology. New York, USA: Springer. 2008. p. 105–20.

8. Chepstow-Lusty AJ, Frogley MR, Bauer BS, Leng MJ, Boessenkool KP, Carcaillet C, et al. Putting the rise of the Inca Empire within a climatic and land management context. Clim Past. 2009, 5: 375–88.

9. Dillehay TD, Kolata AL. Long-term human response to uncertain environmental conditions in the Andes. Proc Natl Acad Sci USA. 2004, 101: 4325–30.

10. Fehren-Schmitz L, Haak W, Maechtle B, Masch F, Llamas B, Tomasto Cagigao E, et al. Climate change underlies global demographic, genetic, and cultural transitions in pre-Columbian southern Peru. Proc Natl Acad Sci USA. 2014, 111: 9443–8.

11. Nielsen AE. Pastoralism and the non-pastoral world in the Late Pre-Columbian history of the Southern Andes (1000-1535). Nomadic Peoples. 2009, 13: 17–35.

12. Aschero CA, Hocsman S. Arqueología de las ocupaciones cazadoras-recolectoras de fines del Holoceno medio de Antofagasta de la Sierra (puna meridional argentina). Chungara. 2011 2011, 43: 393–411.

13. Morales M, Barberena R, Belardi JB, Borrero L, Cortegoso V, Duran V, et al. Reviewing human-environment interactions in arid regions of southern South America during the past 3000 years. Paleogeogr Paleoclimatol Paleoecol. 2009, 281: 283–95.

14. Babot MP. Cazadores-recolectores de los Andes centro-sur y procesamiento vegetal: una discusión desde la puna meridional argentina (ca. 7.000-3.200 años A.P.). Chungara. 2011, 43: 413–32.

15. Planella MT, Scherson R, McRostie V. Sitio El Plomo y nuevos registros de cultígenos iniciales en cazadores del Arcaico IV en Alto Maipo, Chile central. Chungara. 2011, 43: 189–202.

16. Cruz P, Winkel T, Ledru M-P, Bernard C, Egan N, Swingedouw D, et al. Rain-fed agriculture thrived despite climate degradation in the pre-Hispanic arid Andes. Sci Adv. 2017, 3:e1701740.

17. Salminci P, Tchilinguirian P, Lane K. Bordos and boundaries: sustainable agriculture in the high altitude deserts of Northwest Argentina, AD 850-1532. J Anthropol Archaeol. 2014, 2: 189–218.

18. Izquierdo AE, Grau HR. Agriculture adjustment, land-use transition and protected areas in Northwestern Argentina. J Environ Manage. 2009, 90: 858–65.

19. Alcalde JA, Kulemeyer JJ. The Holocene in the South-Eastern region of the Province Jujuy, North-West Argentina. Quatern Int. 1999, 57–8:113–6.

20. Lupo LC, Bianchi MM, Araoz E, Grau R, Lucas C, Kern R, et al. Climate and human impact during the past 2000 years as recorded in the Lagunas de Yala, Jujuy, northwestern Argentina. Quatern Int. 2006, 158: 30–43.

21. Ratto N, Montero C, Hongn F. Environmental instability in western Tinogasta (Catamarca) during the Mid-Holocene and its relation to the regional cultural development. Quatern Int. 2013 12, 307: 58–65.

22. Schittek K, Kock ST, Lücke A, Ohlendorf C, Kulemeyer JJ, Lupo LC, et al. Environmental and climatic history in the NW Argentine Andes (24° S) over the last 2100 years inferred from a high-altitude peatland record. Clim Past. 2015, 11: 2037–76.

23. Tchilinguirian P, Olivera DE. Late quaternary paleoenvironments, south Andean puna (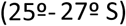), Argentina. In: Pintar E, editor. Hunter-gatherers from a high-elevation desert : people of the Salt Puna : Northwest Argentina. Oxford, UK: Archaeopress. 2014. p. 43–69.

24. Babot P, Hocsman S. Quinoa: a millenary grain in Northern Argentina. In: Selin H, editor. Encyclopaedia of the History of Science, Technology, and Medicine in Non-Western Cultures. Dordrecht, The Netherlands: Springer. 2015.

25. Lopez LM, Recalde AM. The first quinoa (Chenopodium quinoa Willd) macrobotanical remains at Sierras del Norte (Central Argentina) and their implications in pre-Hispanic subsistence practices. J Archaeol Sci Reports. 2016, 8: 426–33.

26. Costa Tartara SM, Manifesto MM, Bramardi SJ, Bertero HD. Genetic structure in cultivated quinoa (Chenopodium quinoa Willd.), a reflection of landscape structure in Northwest Argentina. Conserv Genet. 2012, 13: 1027–38.

27. Zeder MA, Emshwiller E, Smith BD, Bradley DG. Documenting domestication: the intersection of genetics and archaeology. Trends Genet. 2006, 22: 139–55.

28. Nordborg M. Coalescent theory. In: Baldin GJ, Bishop M, C. C, editors. Handbook of Statistical Genetics 3rd edition. Chichester, UK: John Wiley & Sons. 2007. p. 843–77.

29. Tchilinguirian P, Olivera DE. De aguas y tierras: aportes para la reactivación de campos agrícolas arqueológicos en la puna argentina. Rel Soc Argent Antropol. 2000, 24: 99–118.

30. Curti RN, Andrade AJ, Bramardi S, Velasquez B, Bertero HD. Ecogeographic structure of phenotypic diversity in cultivated populations of quinoa from Northwest Argentina. Ann Appl Biol. 2012, 160: 114–25.

31. Lorandi AM. Las rebeliones indígenas. In: Tandeter E, editor. Nueva historia argentina Tomo 2: La sociedad colonial. Buenos Aires, Argentina: Editorial Sudamericana. 2000. p. 285–329.

32. Jellen EN, Kolano BA, Sederberg MC, Bonifacio A, Maughan PJ, Kole C. Chenopodium. In: Kole C, editor. Wild crop relatives: genomic and breeding resources Legume crops and forages. Berlin, Germany: Springer Verlag. 2011. p. 35–61.

33. Quesada MN, Lema C. Los potreros de Antofagasta: trabajo indígena y propiedad (finales del siglo XVIII y comienzos del XIX). Andes (Salta). 2011, 22:ISSN S1668–8090.

34. del Castillo C, Winkel T, Mahy G, Bizoux JP. Genetic structure of quinoa (Chenopodium quinoa Willd.) from the Bolivian altiplano as revealed by RAPD markers. Genet Resour Crop Evol. 2007, 54: 897–905.

35. Bruno MC. A morphological approach to documenting the domestication of Chenopodium in the Andes. In: Zeder MA, Emshwiller E, Smith BD, Bradley DG, editors. Documenting domestication: new archaeological and genetic paradigms. Berkeley, USA: University of California Press. 2006. p. 32–45.

36. Bruno MC, Whitehead WT. Chenopodium cultivation and formative period agriculture at Chiripa, Bolivia. Lat Amer Antiq. 2003, 14: 339–55.

37. Lema VS. Boceto para un esquema: domesticación y agricultura temprana en el Noroeste argentino. Rev Esp Antropol Amer. 2014, 44: 465–94.

38. Kistler L, Shapiro B. Ancient DNA confirms a local origin of domesticated chenopod in eastern North America. J Archaeol Sci. 2011, 38: 3549–54.

39. Arkush E, Stanish C. Interpreting conflict in the ancient Andes - Implications for the archaeology of warfare. Curr Anthropol. 2005 Fb, 46: 3–28.

40. Nielsen AE. Asentamientos, conflicto y cambio social en el altiplano de Lípez (Potosí). Rev Esp Antropol Amer. 2002, 32: 179–205.

41. Martel AR, Aschero CA. Pastores en acción: Imposición iconográfica vs. autonomía temática. In: Nielsen A, Rivolta MC, Selded V, Vázquez MM, Mercolli PH, editors. Producción y circulación prehispánicas de bienes en el sur andino. Córdoba, Argentina: Editorial Brujas. 2007. p. 329–49.

42. Vinton SD, Perry L, Reinhard KJ, Santoro CM, Teixeira-Santos I. Impact of empire expansion on household diet: the Inka in Northern Chile’s Atacama Desert. PLoS ONE. 2009, 4:e8069.

43. Bar-Yosef O. Climatic Fluctuations and Early Farming in West and East Asia. Curr Anthropol. 2011, 52:S175–S93.

44. Zeder MA. The origins of agriculture in the Near East. Curr Anthropol. 2011, 52:S221–S35.

45. Korstanje MA, Cuenya P, Williams VI. Taming the control of chronology in ancient agricultural structures in the Calchaqui Valley, Argentina. Non-traditional data sets. J Archaeol Sci. 2010, 37: 343–9.

46. Etter A, McAlpine C, Possingham HP. Historical patterns and drivers of landscape change in Colombia since 1500: A regionalized spatial approach. Ann Assoc Am Geogr. 2008, 98: 2–23.

47. Morlon P, Martínez ER, Brougère AM. Comprender la agricultura campesina en los Andes Centrales: Perú-Bolivia. Lima, Peru: Centro de Estudios Regionales Andinos Bartolomé de las Casas. 1996.

48. Wachtel N. La vision des vaincus : la conquête espagnole dans le folklore indigène. Annales. 1967, 3: 554–85.

49. Capparelli A, Lema V, Giovannetti M, Raffino R. The introduction of Old World crops (wheat, barley and peach) in Andean Argentina during the 16th century AD: archaeobotanical and ethnohistorical evidence. Veg Hist Archaeobot. 2005, 14: 472–84.

50. Gade DW. Landscape, system, and identity in the post-conquest Andes. Ann Assoc Am Geogr. 1992, 82: 460–77.

51. Jamieson RW, Sayre MB. Barley and identity in the Spanish colonial Audiencia of Quito: Archaeobotany of the 18th century San Blas neighborhood in Riobamba. J Anthropol Archaeol. 2010, 29: 208–18.

52. Parcero-Oubiña C, Fábrega-Álvarez P, Salazar D, Troncoso A, Hayashida F, Pino M, et al. Ground to air and back again: Archaeological prospection to characterize prehispanic agricultural practices in the high-altitude Atacama (Chile). Quatern Int. 2016, 435: 98–113.

53. Belotti López de Medina CR, López Geronazzo L, Otero C. At the feet of the fortress: analysis of Inka period (ca. AD 1430-1536) archaeofaunal assemblages from residential unit 1 (RU1), Pucara de Tilcara (Jujuy, Argentina). PLoS ONE. 2016, 11:e0163766.

54. Gil Montero R, Villalba R, Luckman BH. Tree rings as a surrogate for economic stress : an example from the Puna of Jujuy, Argentina in the 19th century. Dendrochronologia. 2005, 22:141–7. eng.

55. Pannell JR, Charlesworth B. Effects of metapopulation processes on measures of genetic diversity. Phil Trans R Soc Lond B. 2000, 355: 1851.

56. Cramp LJE, Evershed RP, Lavento M, Halinen P, Mannermaa K, Oinonen M, et al. Neolithic dairy farming at the extreme of agriculture in northern Europe. Proc Roy Soc Lond B. 2014, 281: 20140819.

57. Riehl S, Pustovoytov KE, Weippert H, Klett S, Hole F. Drought stress variability in ancient Near Eastern agricultural systems evidenced by delta C-13 in barley grain. Proc Natl Acad Sci USA. 2014, 111: 12348–53.

58. Fraser EDG. Coping with food crises: Lessons from the American Dust Bowl on balancing local food, agro technology, social welfare, and government regulation agendas in food and farming systems. Glob Environ Change-Human Policy Dimens. 2013, 23: 1662–72.

59. Stafford Smith DM, McKeon GM, Watson IW, Henry BK, Stone GS, Hall WB, et al. Learning from episodes of degradation and recovery in variable Australian rangelands. Proc Natl Acad Sci USA. 2007, 104: 20690–5.

60. Pintar E. Continuidades e hiatos ocupacionales durante el holoceno medio en el borde oriental de la puna salada, Antofagasta de la Sierra, Argentina. Chungara. 2014, 46: 51–71.

61. Rodriguez MF, Rugolo de Agrasar ZE, Aschero CA. El uso de las plantas en unidades domésticas del sitio arqueológico Punta de la Peña 4, puna meridional argentina. Chungara. 2006, 38:257–71.

62. Oliszewski N, Martínez JG, Caria MA. Ocupaciones prehispánicas de altura: el caso de Cueva de los Corrales 1 (El Infiernillo, Tafí del Valle, Tucumán). Rel Soc Argent Antropol. 2008, 33: 209–21.

63. Goudet J. HIERFSTAT, a package for R to compute and test hierarchical F-statistics. Mol Ecol Notes. 2005, 5: 184–6.

64. Jombart T. adegenet: a R package for the multivariate analysis of genetic markers. Bioinformatics. 2008, 24: 1403–5.

65. Kamvar ZN, Brooks JC, Grunwald NJ. Novel R tools for analysis of genome-wide population genetic data with emphasis on clonality. Front Genet. 2015, 6: 208.

66. Szpiech ZA, Jakobsson M, Rosenberg NA. ADZE: a rarefaction approach for counting alleles private to combinations of populations. Bioinformatics. 2008, 24: 2498–504.

67. Pritchard JK, Stephens M, Donnelly P. Inference of population structure using multilocus genotype data. Genetics. 2000, 155: 945–59.

68. Pudlo P, Marin JM, Estoup A, Cornuet JM, Gautier M, Robert CP. Reliable ABC model choice via random forests. Bioinformatics. 2016, 32: 859–66.

69. Raynal L, Marin JM, Pudlo P, Ribatet M, Robert CP, Estoup A. ABC random forests for Bayesian parameter inference: arXiv 1605.05537v4: a preprint reviewed and recommended by Peer Community in Evolutionary Biology., 2017.

70. Hocsman S. Producción lítica, variabilidad y cambio en Antofagasta de la Sierra (ca. 5500-2000 AP) [Doctoral Thesis]. La Plata, Argentina: FCNyM-Universidad Nacional de La Plata. 2006.

71. Paabo S, Poinar H, Serre D, Jaenicke-Despres V, Hebler J, Rohland N, et al. Genetic analyses from ancient DNA. Annu Rev Genet. 2004, 38: 645–79.

72. O’Donoghue K, Clapham A, Evershed RP, Brown TA. Remarkable preservation of biomolecules in ancient radish seeds. Proc Roy Soc Lond B. 1996, 263: 541–7.

73. Lia VV, Confalonieri VA, Ratto N, Hernandez JAC, Alzogaray AMM, Poggio L, et al. Microsatellite typing of ancient maize: insights into the history of agriculture in southern South America. Proc Roy Soc Lond B. 2007, 274: 545–54.

74. Wilson AS, Taylor T, Ceruti MC, Chavez JA, Reinhard J, Grimes V, et al. Stable isotope and DNA evidence for ritual sequences in Inca child sacrifice. Proc Natl Acad Sci USA. 2007, 104: 16456–61.

75. Burrieza HP, Sanguinetti A, Michieli CT, Bertero HD, Maldonado S. Death of embryos from 2300-year-old quinoa seeds found in an archaeological site. Plant Sci. 2016, 253: 107–17.

76. Mason SL, Stevens MR, Jellen EN, Bonifacio A, Fairbanks DJ, Coleman CE, et al. Development and use of microsatellite markers for germplasm characterization in quinoa (Chenopodium quinoa Willd.). Crop Sci. 2005, 45: 1618–30.

77. Jarvis DE, Kopp OR, Jellen EN, Mallory MA, Pattee J, Bonifacio A, et al. Simple sequence repeat marker development and genetic mapping in quinoa (Chenopodium quinoa Willd.). J Genet. 2008, 87:39–51.

78. Thompson JD, Higgins DG, Gibson TJ. CLUSTAL W: improving the sensitivity of progressive multiple sequence alignment through sequence weighting, position-specific gap penalties and weight matrix choice. Nucleic Acids Res. 1994, 22: 4673–80.

79. Meyers BC, Tingley SV, Morgante M. Abundance, distribution, and transcriptional activity of repetitive elements in the maize genome. Genome Res. 2001, 11: 1660–76.

80. Hurlbert SH. Nonconcept of species diversity - critique and alternative parameters. Ecology. 1971, 52: 577–86.

81. Weir BS, Cockerham CC. Estimating F-statistics for the analysis of population-structure. Evolution. 1984, 38: 1358–70.

82. Hartl DL, Clark AG. Principles of Population Genetics. Fourth Edition. Sunderland, MA, USA: Sinauer Associates. 2007.

83. David P, Pujol B, Viard F, Castella V, Goudet J. Reliable selfing rate estimates from imperfect population genetic data. Mol Ecol. 2007, 16: 2474–87.

84. Jombart T, Devillard S, Balloux F. Discriminant analysis of principal components: a new method for the analysis of genetically structured populations. Bmc Genetics. 2010, 11: 94.

85. Cornuet J-M, Pudlo P, Veyssier J, Dehne-Garcia A, Gautier M, Leblois R, et al. DIYABC v2.0: a software to make approximate Bayesian computation inferences about population history using single nucleotide polymorphism, DNA sequence and microsatellite data. Bioinformatics. 2014, 30: 1187–9.

86. Marin JM, Raynal L, Pudlo P, Robert CP, Estoup A. abcrf: Approximate Bayesian computation via random forests. R package version 17. 2017.

